# Phase transition and amyloid formation by a viral protein as an additional molecular mechanism of virus-induced cell toxicity

**DOI:** 10.1101/497024

**Authors:** Edoardo Salladini, Claire Debarnot, Vincent Delauzun, Maria Grazia Murrali, Priscila Sutto-Ortiz, Silvia Spinelli, Roberta Pierattelli, Christophe Bignon, Sonia Longhi

**Affiliations:** Aix-Marseille Univ, CNRS, Architecture et Fonction des Macromolécules Biologiques (AFMB), UMR 7257, Marseille, France; Magnetic Resonance Center (CERM), Department of Chemistry “Ugo Schiff” Via Luigi Sacconi 6, 50019 Sesto Fiorentino, Italy

**Author notes:** to whom correspondence should be sent Sonia Longhi, AFMB, UMR 7257 CNRS and Aix-Marseille University, 163, avenue de Luminy, Case 932, 13288 Marseille Cedex 09, France.

**Keywords:** Hendra virus, V protein, intrinsically disordered proteins, amyloids, fibrils, phase transitions

## Abstract

Henipaviruses are severe human pathogens responsible for severe encephalitis. Their V protein is a key player in the evasion of the host innate immune response. We have previously reported a biophysical characterization of the *Henipavirus* V proteins and shown that they interact with DDB1, a cellular protein that is a component of the ubiquitin ligase E3 complex. Here, we serendipitously discovered that the Hendra virus V protein undergoes a liquid-hydrogel phase transition. By combining experimental and bioinformatics approaches, we have identified the V region responsible for this phenomenon. This region (referred to as PNT3), which falls within the long intrinsically disordered region of V, was further investigated using a combination of biophysical and structural approaches. ThioflavinT and Congo red binding assays, together with negative-staining electron microscopy studies, show that this region forms amyloid-like, β-enriched structures. Such structures are also formed in mammal cells transfected to express PNT3. Those cells also exhibit a reduced viability in the presence of a stress agent. Interestingly, mammal cells expressing a rationally designed, non-amyloidogenic PNT3 variant (PNT3^3A^), appear to be much less sensitive to the stress agent, thus enabling the establishment of a link between fibril formation and cell toxicity. The present findings therefore pinpoint a so far never reported possible mechanism of virus-induced cell toxicity.

## Introduction

The Hendra and Nipah viruses (HeV and NiV) are members of the *Paramyxoviridae* family within the *Mononegavirales* order that comprises non-segmented, negative-stranded RNA viruses. NiV and HeV are zoonotic agents responsible for severe encephalitis in humans that have been classified within the *Henipavirus* genus [1]. HeV emerged in 1994 as the etiologic agent of a sudden outbreak of acute respiratory and neurological disease in horses in Brisbane, Australia. HeV infections presently constitute a serious threat to livestock in Australia, where sporadic and lethal transmission to humans has also occurred. After the first cases of human infection in 1998 in Malaysia, NiV has regularly reemerged since 2001 in Bangladesh, with an average case fatality of 70%. The ability of NiV to be transmitted by direct inter-human transmission further extends its potential to cause deadly outbreaks [2-4]. In addition, the discovery of Henipaviruses in other species of bats in West Africa and China [5] underscores the threat that these viruses constitute to human health. The susceptibility of humans, their high pathogenicity, the wide host range and interspecies transmission and the lack of vaccines and therapeutic treatments led to the classification of henipaviruses as biosecurity level 4 (BSL-4) pathogens and as potential bio-terrorism agents.

Like for all *Mononegavirales* members, the genome of henipaviruses is encapsidated by the nucleoprotein (N) within a helical nucleocapsid that serves as the substrate used by the viral polymerase for both transcription and replication. The viral polymerase is a complex made of the large (L) protein and the phosphoprotein (P). The P protein is an essential polymerase cofactor: not only it allows recruitment of L onto the nucleocapsid template but also serves as a chaperon for L [6, 7].

*Henipavirus* P proteins consist in an exceptionally long (> 400 aa), N-terminal disordered region (PNT) [8] and in a C-terminal region of approximately 300 aa containing two structured regions: a coiled-coil domain responsible for P multimerization (PMD) [9-11], and the X domain (XD). The latter adopts a triple α-helical bundle fold [12] and is responsible for interaction with the C-terminal disordered domain (N_TAIL_) of the N protein [13-15]. In paramyxoviruses, the P gene also encodes the V and W proteins that are produced upon addition of either one (protein V) or two (protein W) non-templated guanosines at the editing site of the P messenger. The P, V and W proteins therefore share the PNT region that can be considered as a *bona fide* domain (**Figure 1A**). The C-terminal domain unique to paramyxoviral V proteins is predicted to be structured and to bind zinc [16] and is therefore referred to as zinc-finger domain (ZnFD).

**Figure 1.**
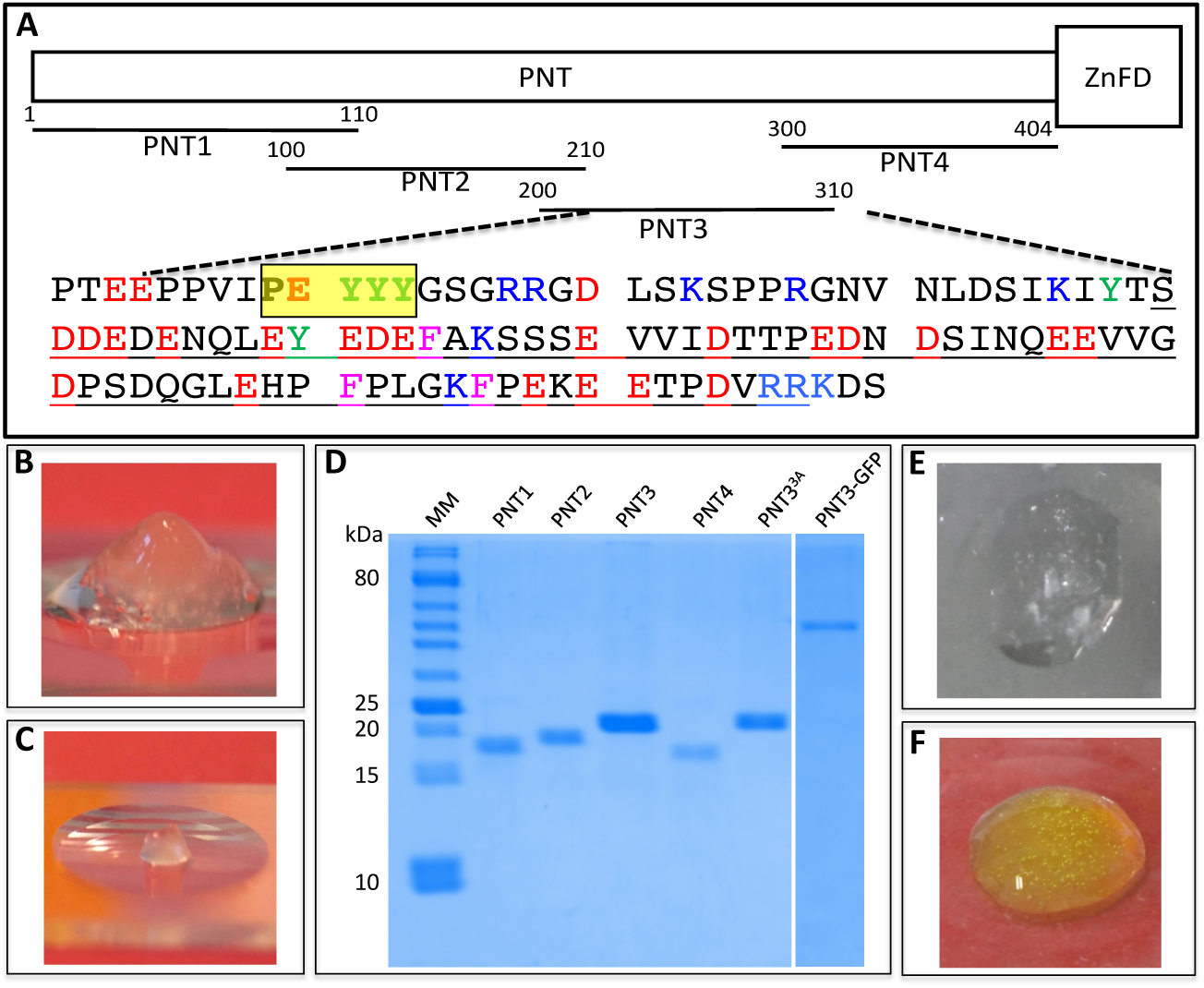
**(A)** Schematic organization of the HeV V protein. The intrinsically disordered P N-Terminal domain (PNT) and the zinc-finger domain (ZnFD) are represented as a narrow and large box respectively. The various PNT fragments herein generated are shown. The amino acid sequence of PNT3 is shown, with the most amyloidogenic region, as predicted by FoldAmyloid, framed in yellow. The low complexity region is underlined. Tyr residues are shown in green, acidic residues in red and basic residues in blue. **(B)** and **(C)** Hydrogels formed by concentrated (200 μM) V **(B)** and PNT **(C)** proteins upon freezing and thawing. **(D)** SDS-PAGE analysis of purified PNT fragments, of PNT3^3A^ and of PNT3-GFP. MM: molecular mass markers. **(E)** and **(F)** Hydrogels formed by PNT3 at 200 μM **(E)** and PNT3-GFP at 1 mM **(F)** upon freezing and thawing.

Paramyxoviral V and W proteins are key players in the evasion of the interferon (IFN) mediated response. They act both by antagonizing IFN signaling and by inhibiting IFN induction [17, 18]. One of the IFN signaling pathways relies on STAT proteins (STAT1 and STAT2) activation and subsequent nuclear translocation. Once imported in the nucleus, they interact with IRF-9 to form the ISGF3 complex that activates the transcription of IFN-stimulated genes whose products inhibit viral replication [17]. The V and W proteins of henipaviruses have an antagonist activity of IFN signaling [17, 19]. They both bind to STAT1 via their PNT domain [20] leading to either inhibition of STAT1 translocation into the nucleus (V) or STAT1 sequestration in the nucleus (W) [20].

Although the *Henipavirus* P protein has an anti-IFN function too, its IFN antagonist activity is moderate compared to V (or W). This observation advocates for a critical role of the C-terminal domain of V and W in the anti-IFN function. That the *Henipavirus* V plays a pivotal role in the evasion of the innate immune response is corroborated by the fact that the Cedar virus (the lastly discovered *Henipavirus* member), which lacks the V protein, induces an IFN response much more pronounced compared to HeV, as well as an asymptomatic infection in animal models [21]. The fact that NiV and HeV are the paramyxoviruses with the highest frequency of P messenger editing and also highly pathogenic comes in further support of a critical role of V in counteracting the innate immune response [18].

Several paramyxoviruses are able to hijack the cellular ubiquitin ligase E3 complex to promote the rapid degradation of STAT proteins. This activity relies on the ability of their V protein to bind to DNA damage-binding protein 1 (DDB1), a component of the ubiquitin ligase E3 complex, and then to recruit STAT proteins onto this complex [22]. In line with this, we have recently reported that *Henipavirus* V proteins interact with DDB1, and have unveiled a critical contribution of their C-terminal domain [23]. Incidentally, we also confirmed that the latter is folded and has a high β content both in isolation and when appended to PNT. Likewise, we have shown that PNT retains its overall disordered nature also in the context of the V protein. This finding rules out the possibility that PNT might adopt a unique conformation in the context of the V protein from which function could arise, and rather argues for a scenario where it is the C-terminal ZnFD that specifically confers to the V protein the ability to counteract the host immune response.

In the course of a further characterization of the *Henipavirus* V proteins, prompted by the fact that they are promising targets for antiviral approaches, we serendipitously discovered that the HeV V protein has the ability to form a hydrogel. In light of the growing number of studies pointing to a critical role of phase separations and transitions mediated by intrinsically disordered proteins (IDPs) and regions (IDRs) in various biological processes [24-26], we have decided to investigate in details this peculiar behavior.

By combining experimental and bioinformatics approaches, we have identified the V region responsible for this phenomenon and have further investigated it using a combination of biophysical and structural approaches. Using Congo Red (CR) and thioflavin T (ThT) binding assays and negative-staining electron microscopy we show that this region forms amyloid-like fibrils. Finally we show that mammal cells transfected to express this region display an increased sensitivity (i.e. increased mortality) towards a stress agent.

## Results

### Liquid-hydrogel transitions by the HeV V protein and identification of the region responsible for this behavior

In view of an in-depth structural characterization of the *Henipavirus* V proteins, we purified large amounts of both NiV and HeV V proteins at a relatively high concentration (i.e. ≥ 10 mg/mL, 200 μM) and stored them at −20°C. Upon thawing them, we noticed that the HeV V protein forms a hydrogel (**Figure 1B**). This liquid-hydrogel transition is irreversible as neither dilution nor boiling can restore the liquid state.

In order to identify the region responsible for this peculiar behavior, we resorted to dividing the V protein in various fragments. Out of the two already available HeV domains (i.e. ZnFD and PNT) [8, 23], only PNT was found to be able to form a hydrogel under conditions similar to those used for HeV V (Figure 1C). We subsequently divided the HeV PNT domain in four overlapping fragments (referred to as PNT1-PNT4) of 110 residues each (**Figure 1A**). The PNT fragments were all purified to homogeneity (**Figure 1D**). PNT3 (aa 200-310) was identified as the only V protein fragment able to form a gel after a freezing/thawing cycle at 200 μM (**Figure 1E**). This ability is also retained in the context of a PNT3-GFP fusion although gel formation was observed at a higher protein concentration (**Figure 1F**).

The ability of PNT3 to form a hydrogel after a freezing/thawing cycle is in line with bioinformatics analyses that unveiled the presence of a low complexity region (enriched in Glu) encompassing residues 240-307. Low complexity domains are indeed known to drive physiologically reversible assembly of IDPs into membrane-free organelles, liquid droplets and hydrogel-like structures [26-30]. In addition, PNT3 contains a stretch of three contiguous tyrosines (aa 210-212 of P), and two non-contiguous tyrosines at positions 237 and 249 **(Figure 1A**). The well-established effect of tyrosines [31] and, more generally, of π-orbital containing residues [32, 33] in promoting phase separation provides an additional conceptual piece to rationalize the behavior of PNT3.

Liquid-hydrogel transitions are thought to be preceded by a step of liquid-liquid phase separation (LLPS) [26, 34] that can be tackled by a wide range of physico-chemical solution studies. We thus first assessed whether PNT3 phase separates and then investigate the conditions under which this phenomenon occurs.

### PNT3 undergoes phase separation

After a 1-hour incubation at room temperature (RT) in the presence of a crowding agent (i.e. 30% PEG300), PNT3 was found to phase separate in the 80 – 240 μM range and to form a hydrogel at 320 μM (see inset in **Figure 2A)**. Phase separation can be quantified by turbidity measurements **(Figure 2A)** that showed that the phenomenon is dependent on both protein and PEG concentration, with a PEG concentration of 20% having no significant impact. We next assessed whether PNT3 retains the ability to aggregate in the absence of a crowding agent and investigated the impact of salt and temperature by monitoring the formation of aggregates. As shown in **Figure 2B**, aggregation occurs in a concentration-dependent manner even in the absence of PEG. Interestingly, salt does not seem to significantly affect the ability of the protein to form aggregates, suggesting that the phenomenon is triggered by the formation of a hydrophobic core. The formation of aggregates is slightly enhanced at 37°C **(Figure 2B)**.

**Figure 2.**
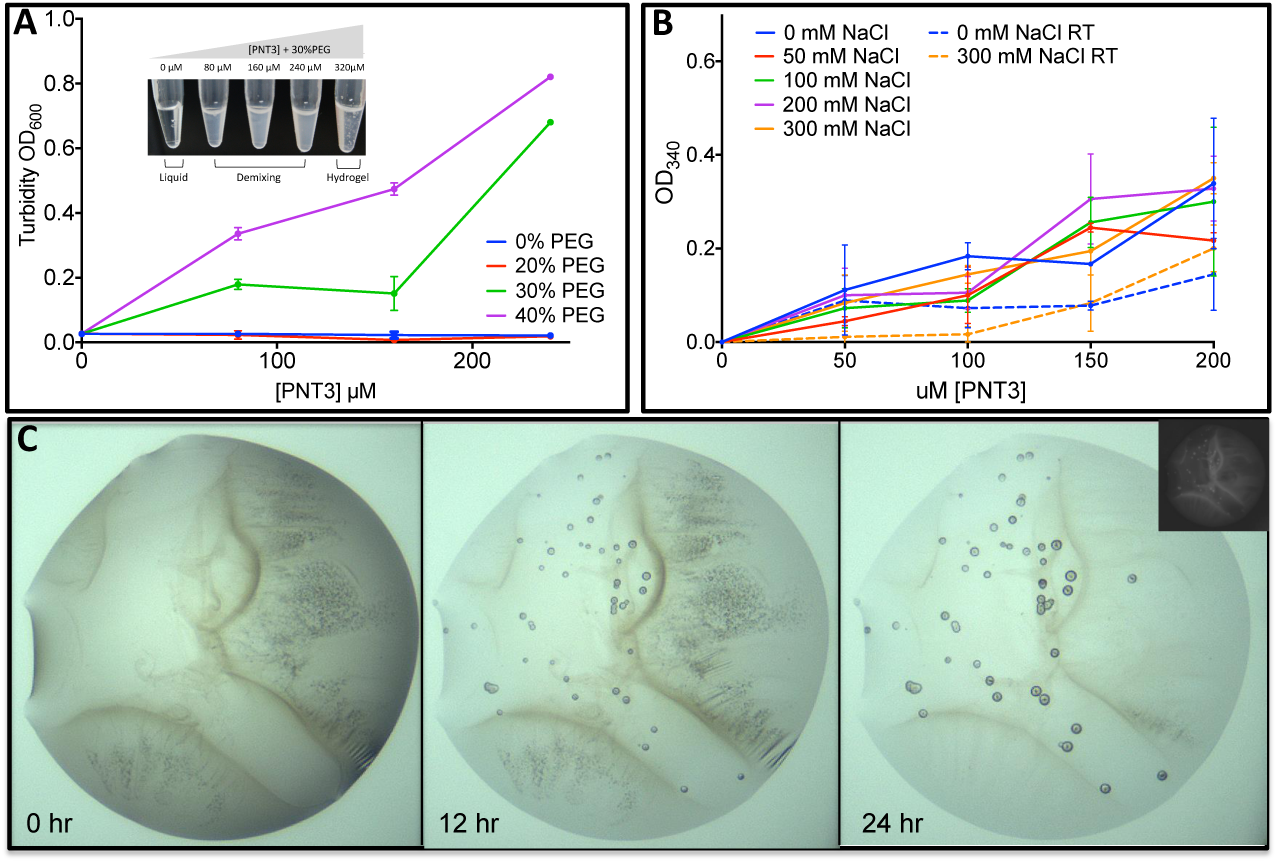
Liquid-liquid phase separation (LLPS) of PNT3. **(A)** Turbidity measurements of PNT3 samples at different concentrations either in the absence or in the presence of increasing PEG 400 concentrations after 1 hour incubation at RT. **(B)** Aggregate formation in PNT3 samples at different concentrations either in the absence or in the presence of increasing concentrations of NaCl after 1 hour incubation at 37°C (continuous lines) or at RT (dotted lines). **(C)** Droplets of 0.3 μl of 250 μM PNT3 in the presence of 10% PEG 4000 at day 0, 1 and 2 of incubation at RT.

A time course analysis of phase separated PNT3, as obtained after incubation at RT of a PNT3 solution at 250 μM supplemented with 10% PEG_4000_, shows the appearance of droplets after 1 day of incubation **(Figure 2C**, central panel). Those droplets become larger between 24 and 48 hours, exhibit a relatively slow dynamics and seem to be rather stable (i.e. all droplets observed after one day can also be observed in the same position after two days). These features suggest a solid-like nature. Indeed, if the droplets were liquid like, Ostwald ripening (i.e. small droplets disappearing in favor of large ones) or coalescence would be expected. Definite answers as to the liquid *versus* solid nature of these droplets in this regard however await fluorescence recovery after photobleaching (FRAP) [35] studies. The proteinaceous nature of those droplets was confirmed under UV light (see inset in **Figure 2C**).

In conclusion, these experiments show that the phase separation undergone by PNT3 is multiparametric, depending on protein concentration, temperature, time and crowding agents.

### PNT3 self-assembles into amyloid-like supramolecular structures

We next sought at shedding light onto the nature of these aggregates. To this end, we analyzed the ability of PNT3 to bind the amyloid-specific dye CR [36]. When the dye is bound to cross β-sheet structures there is hyperchromicity and a red shift of the absorbance maximum. The addition of 25 μM PNT3 promotes a shift in the CR spectrum from 497 nm to 515 nm (**Figure 3A**).

**Figure 3.**
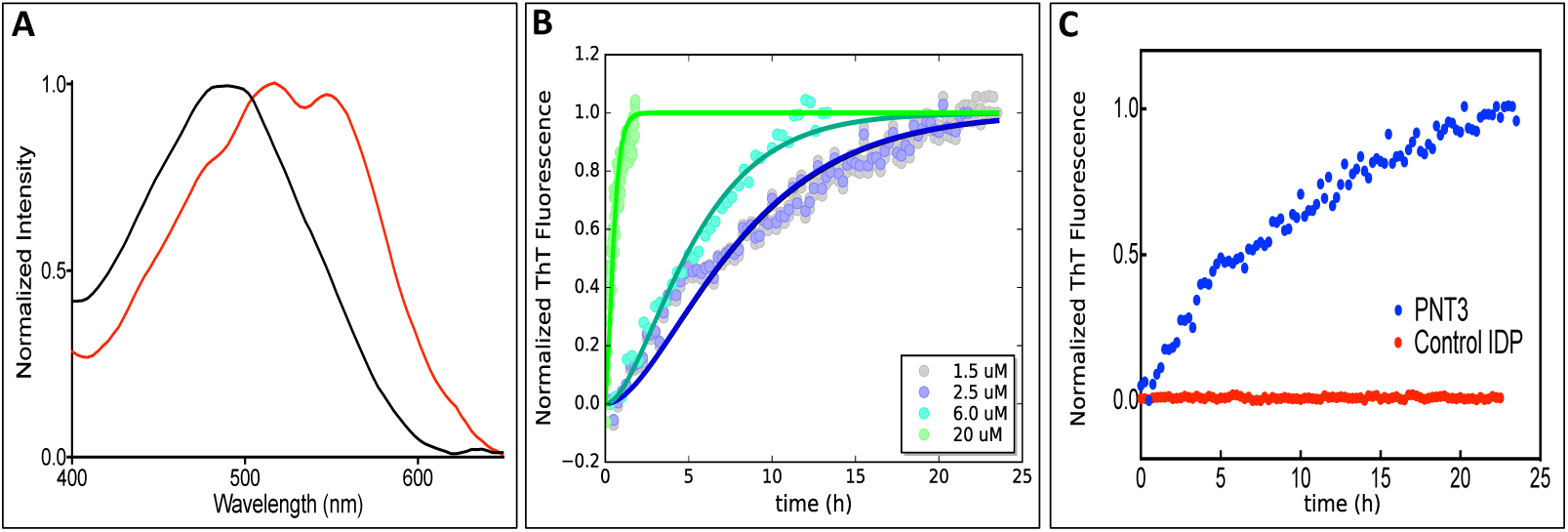
Congo Red and Thiofavin T binding assays. **(A)** Congo red (CR) spectral changes in the absence (black line) or in the presence (red line) of a 25 μM PNT3 sample at after 27 hours of incubation at 37°C. Note the characteristic shift from 497 nm to 515 nm typical of the dye when bound to amyloid-like aggregates. **(B)** Thioflavin T (ThT) binding assays of PNT3 at various concentrations and at various time points of incubation at 37°C. **(C)** ThT binding assays of PNT3 and of a control IDP (measles virus N_TAIL_) at 6 μM. The fluorescence was measured at various time points of incubation at 37°C.

This observation provides the first hint suggesting that PNT3 can form β-enriched/amyloid-like structures. To quantify this phenomenon, we carried out binding assays with ThT, another well-known amyloid-specific dye [37]. Binding of ThT to PNT3 was found to induce a large enhancement in the intensity of the ThT fluorescence emission spectrum in a time- and concentration-dependent manner **(Figures 3B** and **3C)**. In particular, while PNT3 samples at 1.5 and 2.5 μM give rise to a very similar kinetics, a significantly faster kinetics is observed at 6.0 μM **(Figures 3B)**. When the protein concentration is increased to 20 μM the kinetics is much faster, with the plateau being reached already after 2 hours **(Figures 3B)**.

The ability of PNT3 to bind both CR and ThT is consistent with bioinformatics analyses carried out using the FoldAmyloid predictor (http://bioinfo.protres.ru/fold-amyloid/) that identified a motif (aa 208-212) containing three contiguous tyrosines as the most amyloidogenic region (see **Figure 1A**). This is also in line with the lack of a significant effect of salt on PNT3 aggregation **(Figure 2B)**, indicative of the presence of a hydrophobic core. These results therefore argue for the ability of PNT3 to form amyloid-like assembles.

### Far-UV circular dichroism studies of PNT3

To achieve additional insights on the secondary structure content of PNT3 before and after phase separation, we recorded the far-UV CD spectra of PNT3 under different conditions **(Figure 4A**). The CD spectrum of a freshly purified sample of PNT3 is typical of an IDP, as illustrated by the very pronounced negative peak at 200 nm. After incubation at 37 °C for ≥ 16 hours, i.e. a condition where we could document fibril formations in ThT and CR binding assays (see **Figure 3)**, a dramatic decrease in the signal was observed. This phenomenon likely arises from the formation of fibrils that cannot be crossed by the polarized light, and was also observed in the case of phase separated Pro-Arg dipeptide repeats [29]. Interestingly, the CD spectrum of PNT3 hydrogel droplets as obtained following a freezing/thawing cycle shows a disorder to β transition, as judged from the appearance of a peak at 215 nm **(Figure 4A**). Collectively these data support the conclusion that PNT3 forms β-enriched fibrils in phase-separated and hydrogels droplets.

**Figure 4.**
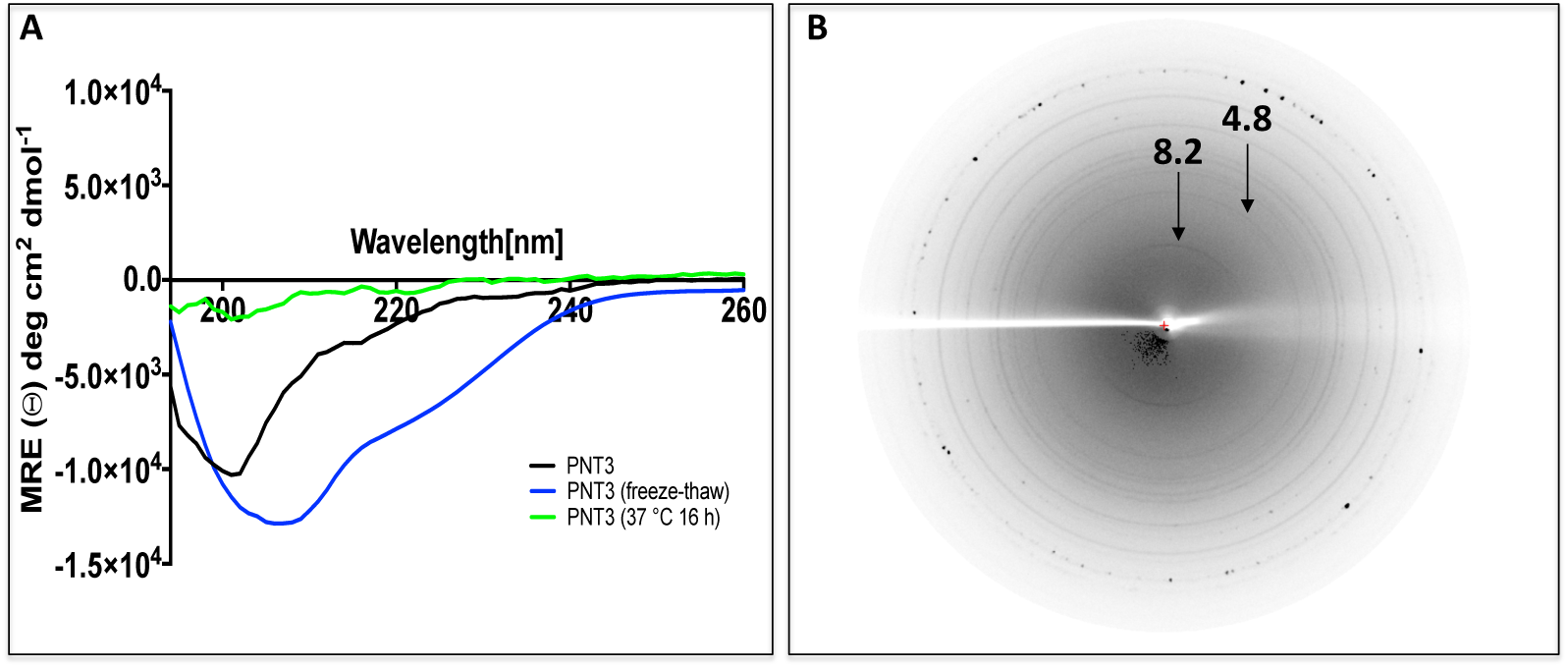
**(A)** Far-UV circular dichroism (CD) studies of PNT3. The spectra were recorded in 10 mM sodium phosphate pH 7 at 20°C. Protein concentration was 0.1 mg/mL (6 μM). The spectrum shown in black corresponds to a freshly purified PNT3 sample recorded immediately after elution from the SEC column. The spectrum shown in green corresponds to a sample incubated at 37°C over night. The spectrum shown in blue corresponds to a sample frozen and thawed. Note the shift from 200 nm to 215 nm of the negative peak in the gelified sample and the dramatic decrease in the signal for the sample incubated over night at 37°C. **(B)** X-ray fiber diffraction pattern of a polymerized PNT3 sample (see Materials and Methods). Prominent, circular X-ray reflections were observed at 4.8 Å and 8.2 Å, which is typical of cross-β structure. The spots at high resolution reflect the diffraction of salt crystals.

### X-ray diffraction studies of PNT3

To assess the presence of cross-β properties in the sample after the ThT assay we employed X-ray diffraction. Fiber diffraction allows visualizing the distances between hydrogen-bonded β-strands (4.7-8 Å diffraction signal on the meridian of the pattern) and the distance that occur from association of the sheets (10-12 Å on the equator, depending on the size of side chains). If the fibers are not aligned on a plane, the diffraction of those distances arises in every direction and appears as a circular intensity, otherwise it is possible to discriminate between meridian and equator axes [38-40]. As shown in **Figure 4B**, the resulting diffraction pattern shows circular reflections at 4.8 Å. There is also a slightly asymmetrical diffraction pattern at 8.2 Å that indicates the equator and suggests a partial alignment of the fibres. These reflections are prototypic of cross-β structure [38-40].

In combination with the ThT binding assays, the X-ray diffraction pattern provides strong evidence of the presence of amyloid-like polymers as the structural basis of phase-separated PNT3.

### Small-angle X-ray scattering studies of PNT3

In view of achieving a better description of the evolution of the conformational properties of PNT3 over time, we carried out SAXS studies. Synchrotron SAXS data were collected from a PNT3 sample at two different concentrations and at different times of incubation (from 0 to 630 min) at 37°C **(Figure 5A** and **Supplementary Figure S1A)**. Overall, the resulting scattering curves show an increase in intensities in the small-angle region with increasing incubation time **(Figure 5A** and **Supplementary Figure S1A)** consistent with an evolution from a monomeric to an oligomeric and/or fibrillar state. We followed this evolution using three parameters, namely the extrapolated forward scattering (I(0)), which is proportional to the molecular mass of the scatterer, the calculated radius of gyration (R_g_) and the maximal internal dimension (D_max_) **(Supplementary Table S1)**.

**Figure 5.**
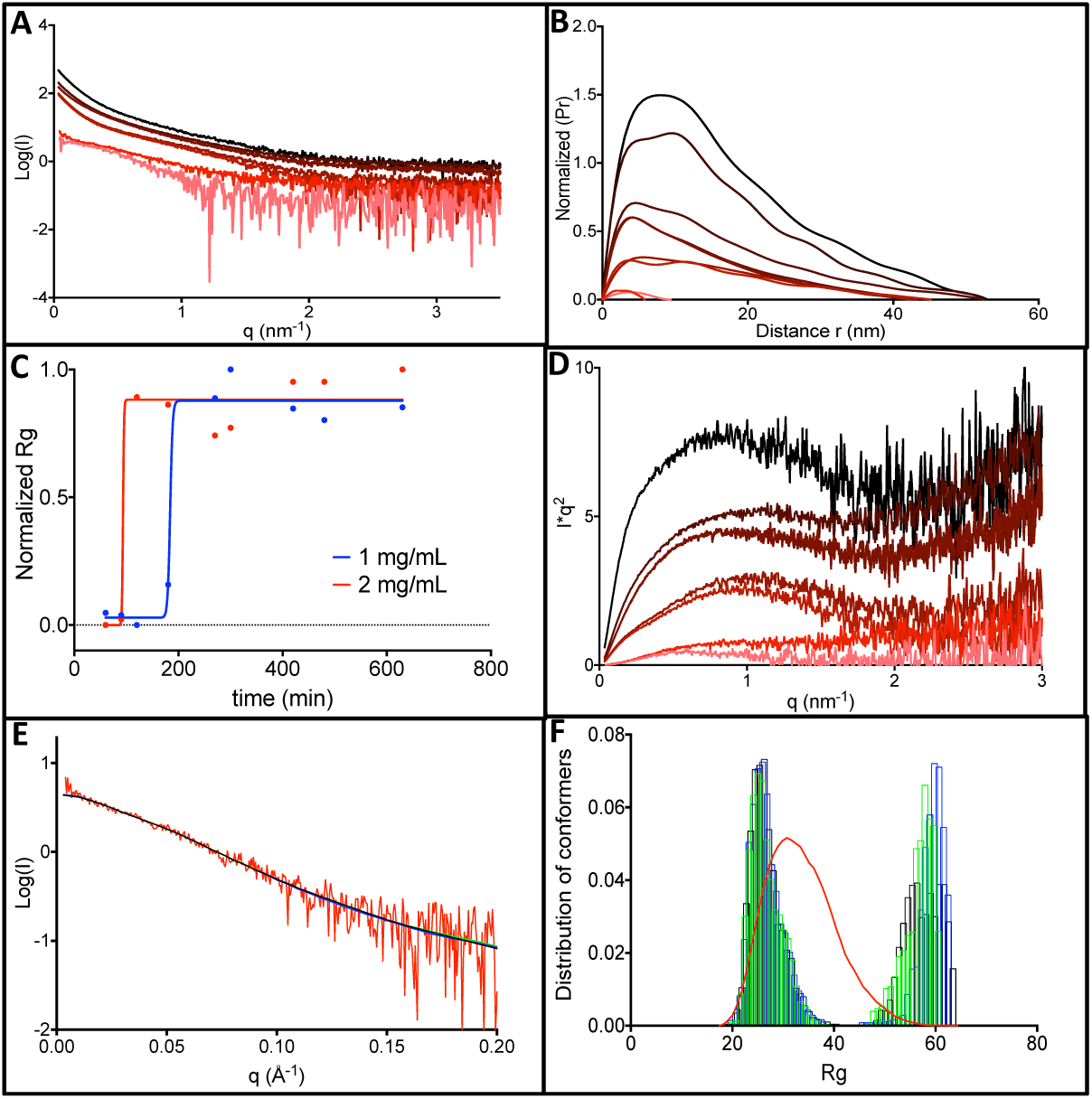
Small-angle X-ray scattering studies of PNT3. **(A)** Scattering curves as obtained from a sample of PNT3 at 2 mg mL^−1^ at various times of incubation at 37°C. The curves are represented with a color gradient ranging from pink to black with increasing incubation times. **(B)** Pair distance distribution, P(r), as obtained at 2 mg mL^−1^. **(C)** Normalized R_g_ as a function of time for the two protein concentrations herein studied. Data were fitted to an asymmetric sigmoid using the Prism software. **(D)** Kratky plot of the SAXS data obtained at 2 mg mL^−1^. The same color code as in panel A was used. **(E)** Experimental scattering curve of the protein at 2 mg mL^−1^ (red) and EOM fits as obtained by three independent runs (black, blue and green). **(F)** R_g_ distribution of the randomly generated conformers by EOM without constraints (red) and of the three sub-ensembles generated by three independent EOM runs (black, blue and green). Note the asymmetrical and bimodal R_g_ distribution.

Both D_max_ **(Figure 5B** and **Supplementary Figure S1B)** and R_g_ **(Supplementary Table S1)** increase with increasing incubation times though in a non-linear manner. In the case of the sample at 1 mg/mL, the monomeric species (calculated molecular mass of 16 kDa) persists until 120 min, while at 2 mg/mL the monomeric species (calculated molecular mass of 16 kDa) can no longer be detected after 90 min **(Supplementary Table S1)**. The kinetics of formation of oligomers and/or fibrils is thus accelerated at higher protein concentrations, in agreement with results from ThT binding experiments (see **Figure 3C)**. This concentration-dependence of fibril formation can be better appreciated in **Figure 5C**, where the R_g_ is plotted as a function of incubation time. The R_g_ value of the monomeric species (32.6 ± 1 Å) is in perfect agreement with the value expected for an IDP of the same size (32.6 Å) according to Flory’s equation [41]. The calculated initial and final values of R_g_, D_max_ and molecular mass **(Figure 5C** and **Supplementary Table S1)** are very similar at the two protein concentrations. The calculated molecular mass after 630 min (see **Supplementary Table S1)** is consistent with an oligomeric/fibrillar species made of ~100 PNT3 monomers.

For both protein concentrations, the Kratky plots reveal an overall gain of content in ordered structure with increasing time, as judged from the appearance of a progressively more pronounced maximum **(Figure 5D** and **Supplementary Figure S1C)**. This increase in ordered structure is consistent with the formation of fibrillar species.

We then investigated the distribution of conformations of the monomeric species at the highest protein concentration using the program suite EOM 2.0 [42]. From an initial pool of 10,000 random conformations, EOM selects a sub-ensemble of conformers that collectively reproduce the experimental SAXS data and represent the distribution of structures adopted by the protein in solution. The average SAXS scattering curves back-calculated from the selected sub-ensembles, as provided by three independent EOM runs, reproduce correctly the experimental curve **(Figure 5E)**. The *R_g_* distribution of the selected sub-ensembles, as obtained from the three independent EOM runs, are asymmetrical and bimodal, with two peaks centered at 25 Å and 60 Å **(Figure 5F)**. The similarity of the R_g_ distributions obtained in different EOM runs attests of the reproducibility of the selection process and hence the reliability of the inferred conformational information.

These data indicate that the scattering curve of the monomeric species of PNT3 does not reflect a randomly distributed ensemble of conformations and *R_g_* distributions and rather the existence of two distinct sub-populations differing in compaction. It is conceivable that the more compact form possesses a higher content in transiently populated secondary structures that imparts a higher propensity to undergo the disorder-to-order transition that accompanies the formation of β-enriched oligomers.

### Nuclear magnetic resonance (NMR) and negative-staining transmission electron microscopy (TEM) studies of PNT3

We next carried out negative-staining TEM and heteronuclear NMR studies to directly document fibril formation as well as any possible significant concomitant structural transition. We thus recorded the ^1^H-^15^N HSQC spectra of a uniformly labeled PNT3 sample after various incubation times at 37°C. As shown in **Figure 6**, a time-course analysis of the sample did not reveal any significant chemical shift displacement. Rather, an overall reduction in the peak intensities was observed with increasing incubation times (see also **Supplementary Figure S2)**. A concomitant analysis with negative staining TEM unambiguously showed the progressive formation of amyloid-like fibrils. While those fibrils become progressively more abundant over time, their diameter (12-17 nm) seemingly remains unvaried. The observed reduction in peak intensities in NMR studies is akin to that observed in CD studies (see **Figure 4A)** and is attributable to the formation of fibrillar species that are not soluble and hence no longer contribute to solution-state NMR signals.

**Figure 6.**
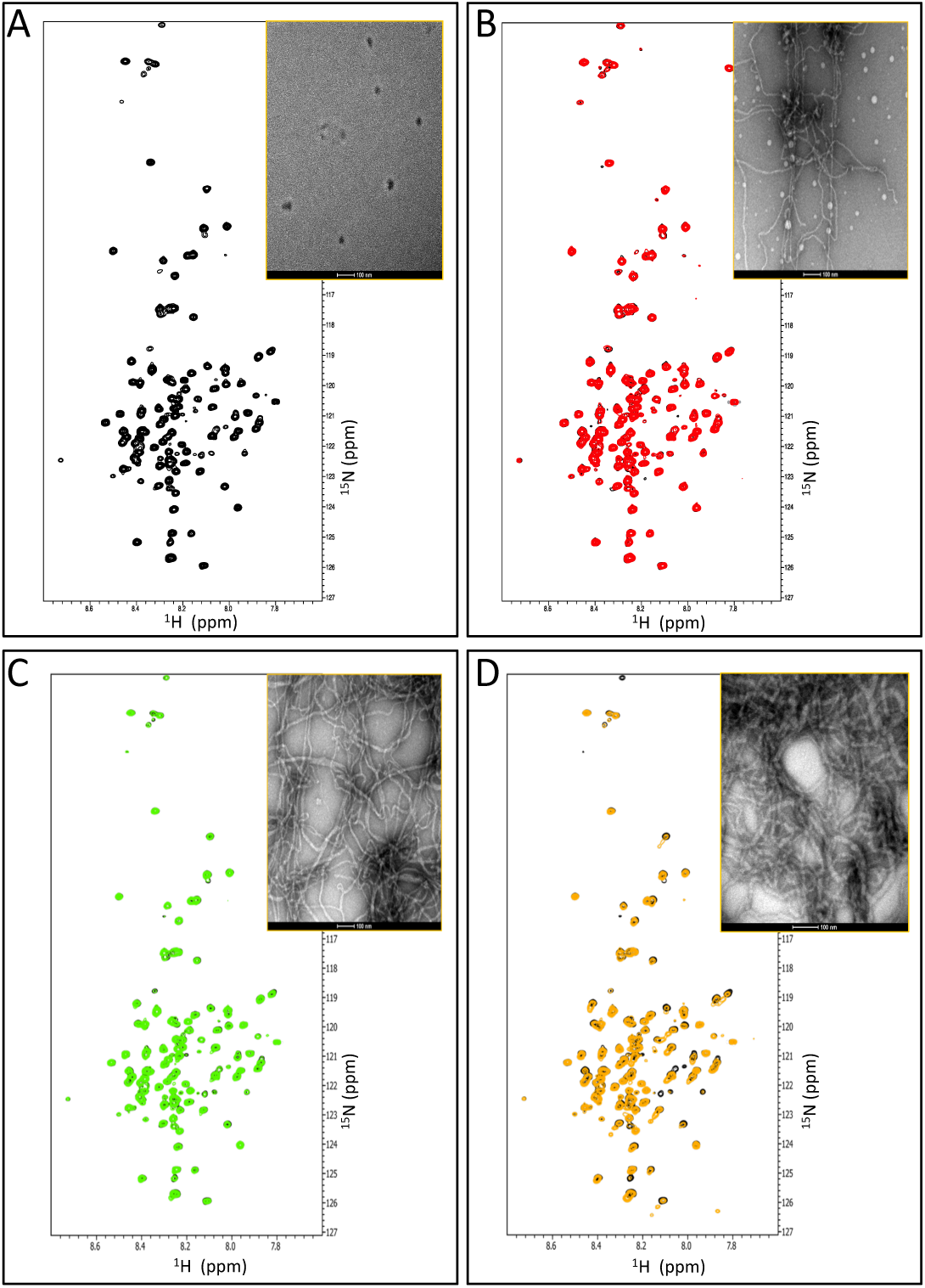
^1^H-^15^N HSQC spectra of a PNT3 sample at 100 μM after 0 **(A)**, 8 **(B)**, 27 **(C)** and 56 **(D)** hours of incubation at 37°C. The HSQC spectrum of the sample at time 0 is shown in black in all panels. The spectra were recorded at 310 K. Insets: negative-staining TEM micrographs of a PNT3 sample at 200 μM incubated at 37°C for 0, 8, 27 and 56 hours.

### Fragility of PNT3 amyloid-like fibers

Extreme stability is a hallmark of pathogenic amyloid fibers (see [43, 44] and references therein cited). In addition, yeast ultra-stable amyloids, derived from the low complexity sequences associated with transcription factors and RNA-binding proteins, have also been described [45]. These prion-like amyloid fibers share a common insensitivity to the solubilizing effects of SDS. In order to investigate the SDS sensitivity of PNT3 fibers, heavily polymerized preparations of PNT3 (i.e. 100 μM after 57 hours of incubation at 37°C) were filtrated through a membrane allowing passage of monomeric proteins but not of fibers. As shown in **Figure 7A**, a very small amount of protein was found to pass through the filter when the fibers were diluted in standard buffer and filtrated immediately. By contrast, following incubation in the presence of 2% SDS at 37° C for 10 min, the amount of UV-adsorbing material passing into the filtrate increases. These results suggest that the fibers were at least partly depolymerized into monomers by SDS treatment. This was further supported by TEM studies that showed that in the presence of 5 mM SDS fibers disappear **(Figure 7B)**.

**Figure 7.**
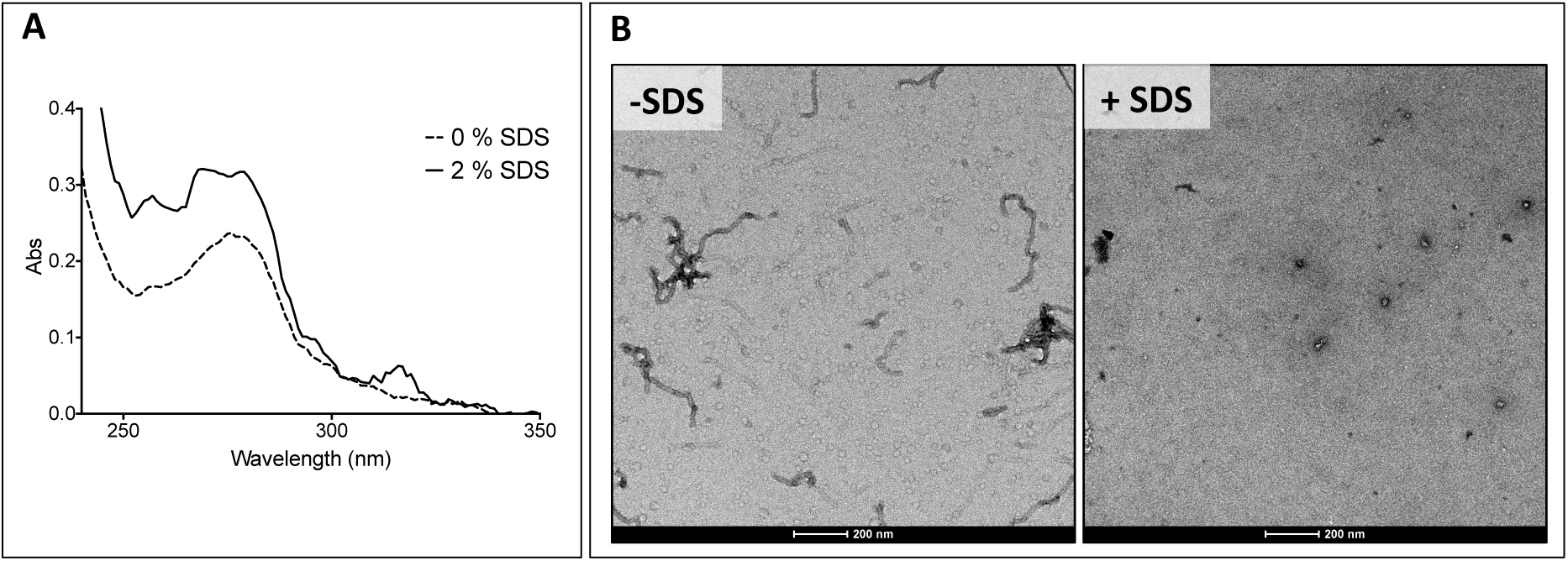
**(A)** UV adsorption of the filtrate of a sample of PNT3 at 100 μM after 56 hours of incubation at 37°C before (dashed line) and after (continuous line) exposure to 2% SDS and 10 min incubation at 37 C. Exposure of SDS enables more material to pass through the filter. **(B)** Micrographs of a PNT3 sample at 200 μM after 56 hours of incubation at 37°C either in the absence or in the presence of 5 mM SDS.

Indeed, analysis of up to 10 grid squares revealed the presence of as few as two small fibers all over. Thus, in the presence of SDS, PNT3 fibrils show a low stability, in line with previous observations on stress granule proteins [46], and on α-synuclein, tau and Aβ42 fibrils [47].

Therefore, although PNT3 fibers share morphological similarities and the presence of cross-β structure properties, they appear to be more fragile than the prion-like fibers broadly described in the literature.

### The heat-shock protein 70 delays the fibrillation process of PNT3

In light of previous studies that documented the ability of chaperons, and in particular of the major inducible heat shock protein 70 (hsp70), to inhibit or delay fibril formation by prion-like proteins [48-52], we decided to ascertain whether human hsp70 has an impact on the fibrillation process of PNT3.

As shown in **Figure 8A**, ThT binding assays showed that hsp70 lowers the rate with which fibrillar species are formed. This observation is mirrored by TEM studies that showed that in the presence of hsp70 the formation of fibrils is hampered **(Figure 8B)**.

**Figure 8.**
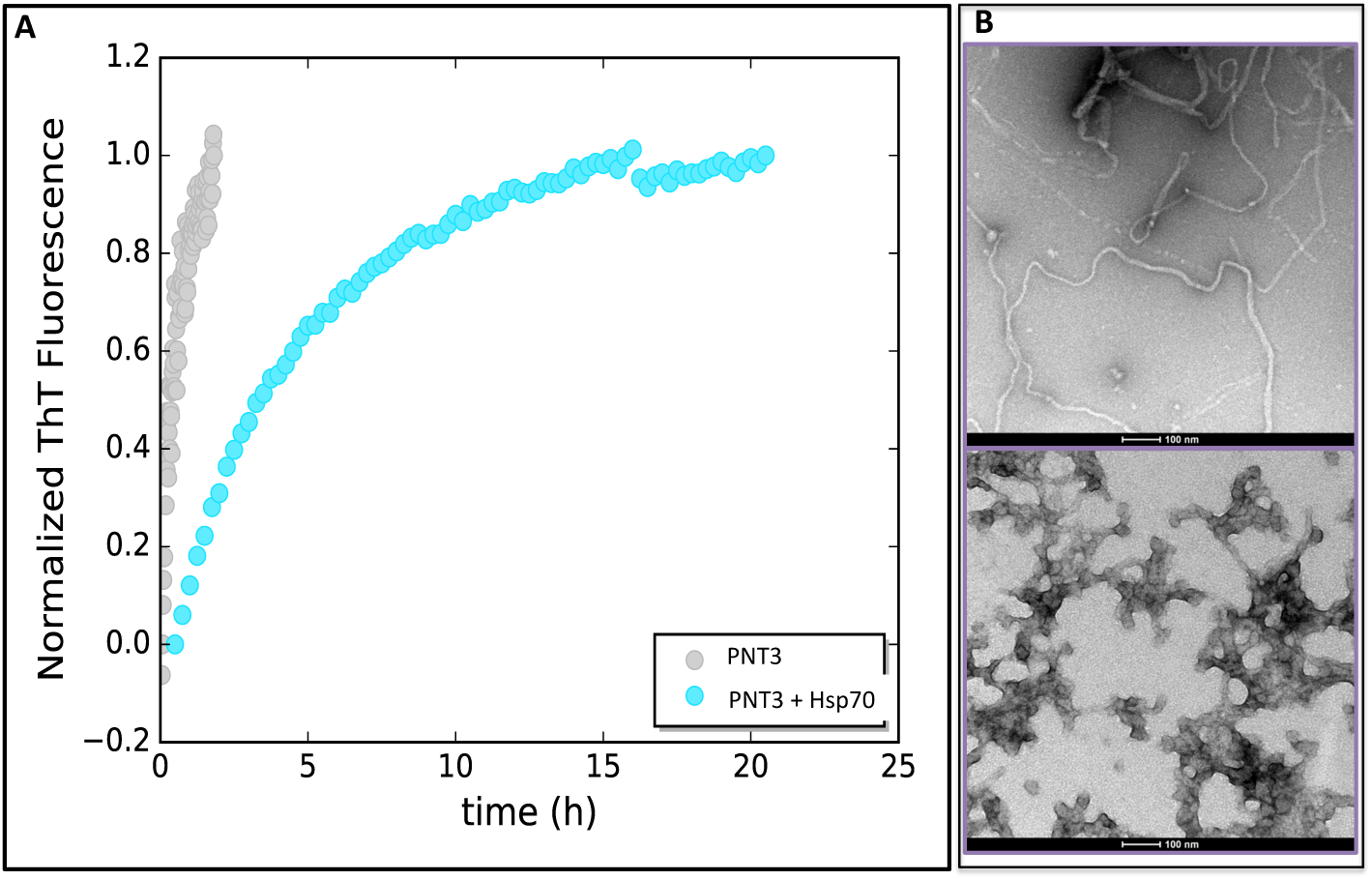
**(A)** ThT binding assays of a PNT3 sample at 20 μM either in the absence or in the presence of a two-fold molar excess of hsp70. The fluorescence was measured at various time points of incubation at 37°C. **(B)** TEM micrographs of a PNT3 sample at 100 μM incubated at 37°C for 27 hours either in the absence (top) or in the presence (bottom) of a two-fold molar excess of hsp70.

### Rational design of a PNT3 variant with a hampered ability to form amyloid-like fibrils

In light of the well-documented role of Tyr residues in LLPS [31, 32], we reasoned that the YYY motif occurring at residues 210-212 of the V protein (see **Figure 1A)** might be responsible for the ability of PNT3 to phase separate and to form amyloid-like fibrils. We thus conceived and generated a PNT3 variant (referred to as PNT3^3A^) in which the three contiguous tyrosine residues were replaced with alanines. The purified PNT3^3A^ variant **(Figure 1B**) has lost the ability to bind CR and ThT (**Figure 9A, B**). The inability of the variant to form amyloid-like fibrils was also confirmed by TEM studies **(Figure 9C)**.

**Figure 9.**
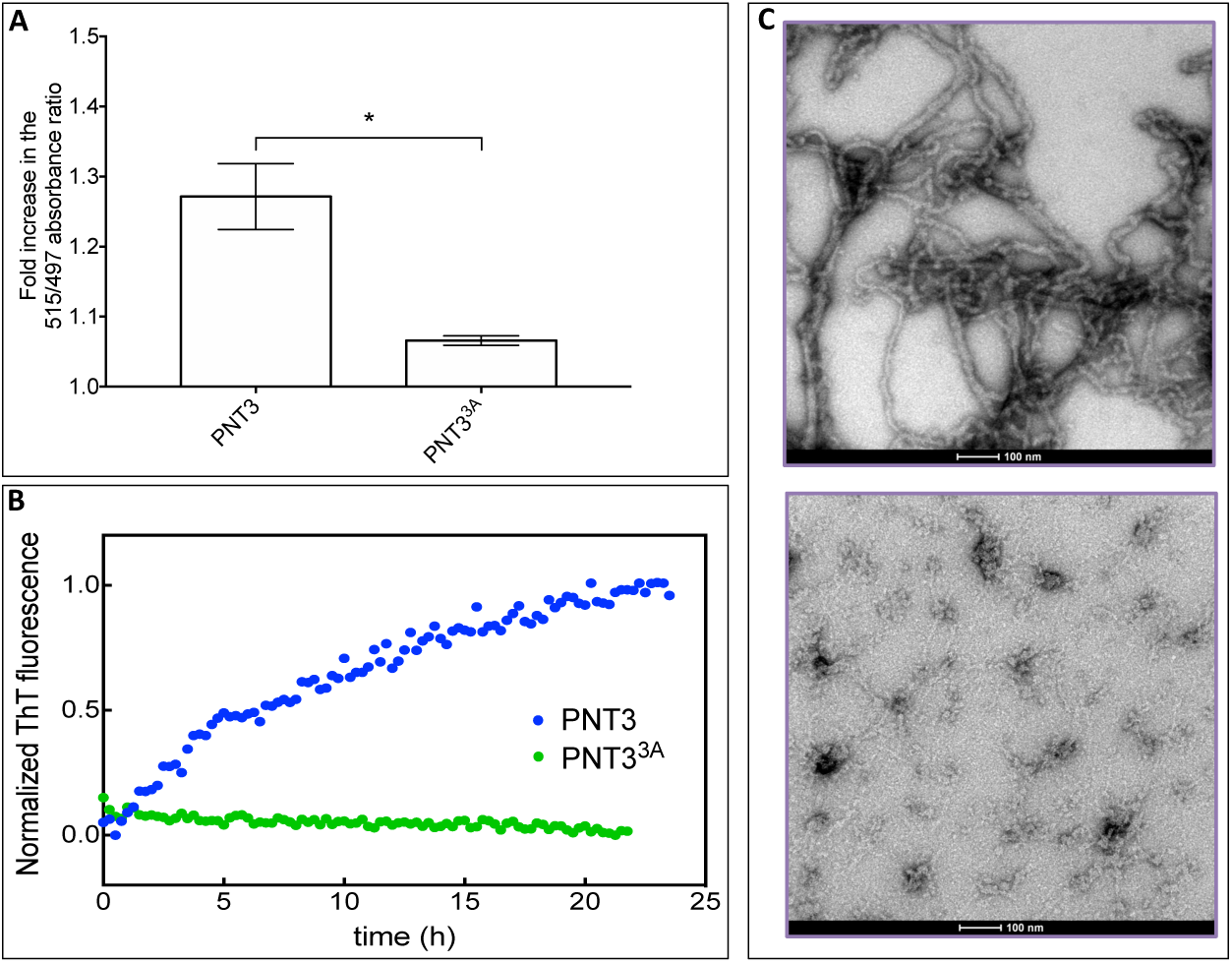
**(A)** CR binding by PNT3 and PNT3^3A^. Fold increase in the ratio between the absorbance at 515 and at 497 nm, with respect to a sample containing CR alone, of a PNT3 or a PNT3^3A^ sample at 25 μM after 27 hours of incubation at 37°C. The error bar corresponds to the standard deviation, with n=3. The asterisk denotes a statistically significant difference (p < 0.001) (T Student’s test). (B) ThT binding assays of PNT3 and of PNT3^3A^ at 6 μM. The fluorescence was measured at various time points of incubation at 37°C. **(C)** TEM micrographs of PNT3 (top) and PNT3^3A^ (bottom) at 200 μM and after a 56-hours incubation at 37°C.

### Functional impact of PNT3 in mammal cells

In an attempt at unraveling the functional impact of the ability of PNT3 to phase separate, we carried out transfection experiments in mammal cells. CR staining of cells transfected to express PNT3 and PNT3-GFP showed that the proteins retain the ability to form amyloid-like fibrils also in the context of transfected cells **(Figure 10A)**. Conversely, and as expected, CR fails to stain non-transfected cells **(Figure 10A)**.

**Figure 10.**
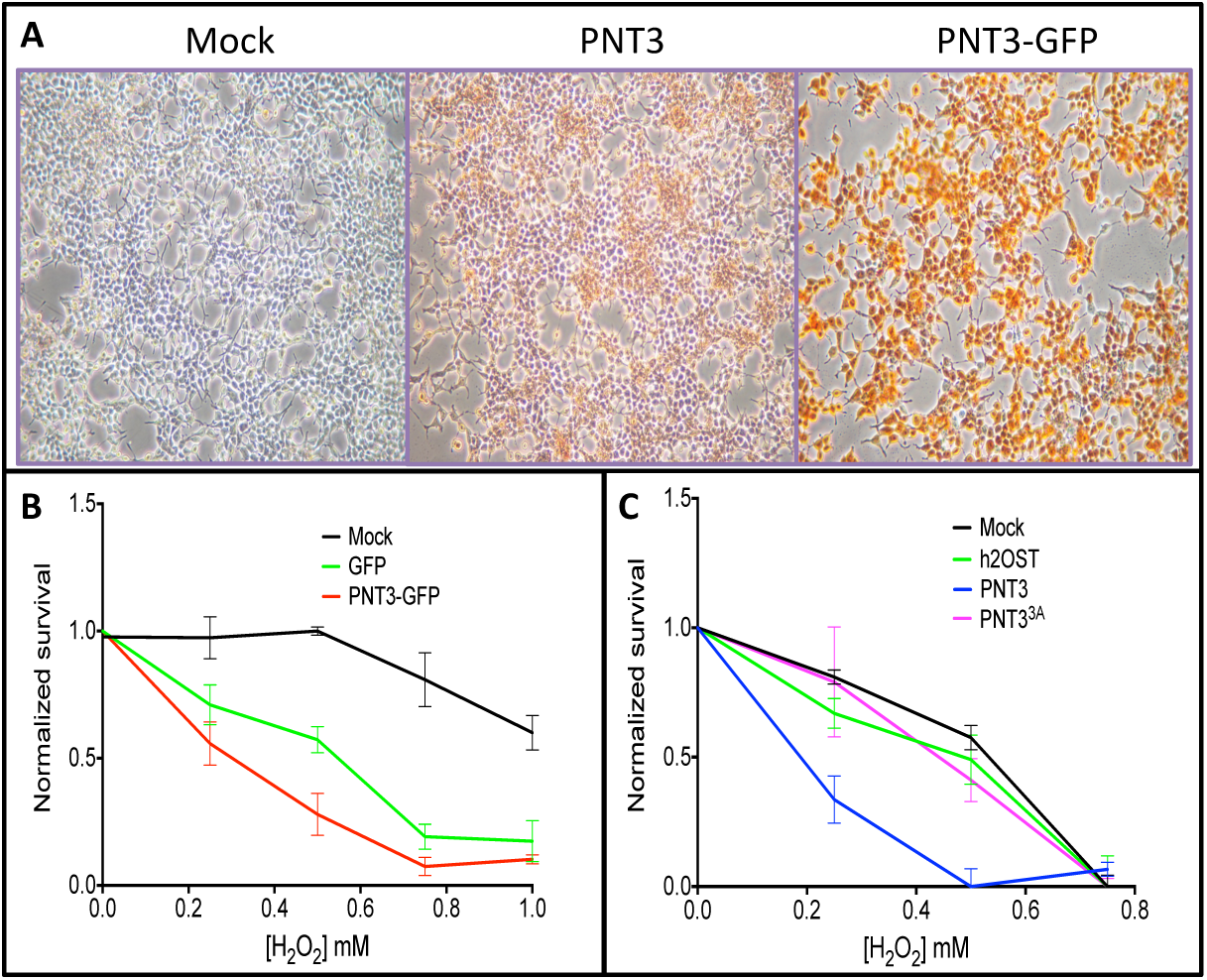
(**A**) CR staining of HEK cells non transfected or transfected to express PNT3 or PNT3-GFP. Note the red staining in the case of PNT3 and PNT3-GFP. **(B)** Viability assays of HEK cells non transfected or transfected to express GFP or PNT3-GFP after 6 hours of incubation at 37°C in the presence of increasing H_2_O_2_ concentrations. Shown are the mean values and s.d. as obtained from a triplicate. **(C)** Viability assays of HEK cells non transfected or transfected to express PNT3 or PNT3^3A^ after 6 hours of incubation at 37°C in the presence of increasing H_2_O_2_ concentrations. Shown are the mean values and s.d. as obtained from a duplicate.

Next we sought at assessing any possible toxic effect related to the formation of amyloid-like fibrils by PNT3. To this end, we first measured cell viability in the presence of a stress agent (i.e. H_2_O_2_). As expected, transfected cells are more sensitive to a stress agent than non-transfected cells **(Figure 10B)**. Interestingly, PNT3-GFP-transfected cells have a lower viability compared to GFP-transfected cells **(Figure 10B)**. Notably, the expression level of PNT3-GFP is 20 times lower than that of GFP alone (as judged from fluorescence measurements, see **Supplementary Figure S3A)** thus ruling out the possibility that the lower toxicity of GFP may arise from a lower expression level. Results therefore advocate for a higher toxicity of PNT3-GFP, with the toxic effect of PNT3 being probably even underestimated.

We next repeated these experiments with cells transfected to express either PNT3 or PNT3^3A^, along with an irrelevant, control protein (i.e. 2OST). As shown in **Figure 10C**, the viability of cells expressing PNT3 in the presence of H_2_O_2_ is much lower compared to that of cells expressing PNT3^3A^, the latter exhibiting a behavior close to that of non-transfected cells and of cells expressing the control protein. The possibility that the lack of toxicity of PNT3^3A^ may arise from a much lower (or lack of) expression in transfected HEK cells was checked and ruled out **(Supplementary Figure S3B)**.

Results advocate for a scenario where the substitution of the triple Tyr motif, which converts PNT3 into a non-amyloidogenic species (see **Figure 9)**, also abrogates toxicity. These findings, especially if also corroborated by future CR staining experiments of PNT3^3A^-transfected cells, allow establishing a link between cell toxicity and ability of the protein to aggregate into amyloid-like structures.

## Discussion

In this paper we showed that the HeV V protein undergoes a liquid to hydrogel transition and mapped the region responsible for this behavior to the long intrinsically disordered region of V and, specifically, to residues 200-310 (PNT3). Using a combination of biophysical and structural approaches, we showed that *in vitro* PNT3 undergoes phase separation and forms amyloid-like, β-enriched structures. Noteworthy, such structures are also formed in mammal cells transfected to express PNT3. Those cells also exhibit a reduced viability in the presence of a stress agent. Interestingly, mammal cells expressing a rationally designed, non-amyloidogenic PNT3 variant (PNT3^3A^), appear to be much less sensitive to the stress agent, thus enabling the establishment of a link between fibril formation and cell toxicity.

In *Mononegavirales*, transcription and replication takes place in viral factories, e.g. cytoplasmic inclusions where the local concentration of viral proteins is increased thereby facilitating viral replication [53-58]. These viral factories, which can be either double-membrane [59] or membrane-less organelles [60], also likely prevent the activation of host innate immunity by shielding viral proteins and thus impairing their interaction with cellular antiviral proteins. It is conceivable that the ability of PNT3 to phase separate may be functionally coupled to the formation of viral factories. This however, would not explain the observed toxicity *per se* of PNT3 fibrils in mammal cells. Noteworthy, recent data suggest that phase transitions of proteins into liquid or hydrogel states could underlie pathological protein aggregation associated with neurodegenerative diseases. In particular, in the case of amyotrophic lateral sclerosis-associated protein FUS (fused in sarcoma) it has been proposed that the toxicity of irreversible FUS assemblies [61] may arise either from the fact that amyloid aggregates are intrinsically toxic for eukaryotic cells, or from the appearance of different density states in the cytoplasm that is the cause of a physiological dysfunction [62]. In this second scenario, the new membrane-less organelles could perturb cellular functions through either sequestration of particular proteins that are thus subtracted from their natural pathways or just by changing the dynamics inside the cells [63].

It is therefore tempting to speculate that the ability of HeV P/V proteins to form amyloids may constitute at least one of the possible molecular mechanisms underlying the pathogenicity of this virus and, possibly, also of its ability to cause encephalitis. The present findings therefore highlight a so far never reported possible mechanism of virus-induced cell toxicity. Definite conclusions however require additional experiments aimed at assessing the presence of amyloid-like fibrils in infected cells. Indeed, the possibility that amyloid formation in transfected cells may arise by a much higher expression level compared to that occurring in the natural context, cannot be ruled out.

To the best of our knowledge, the ability of a viral protein to form amyloids has been reported before only in the case of human papilloma virus (HPV)-16 E7 protein (i.e. the major oncoprotein of HPV). HPV E7 was indeed shown to be able self-assemble into defined spherical oligomers with amyloid-like properties [64, 65], although the possible functional implications of this phenomenon were only discussed in terms of the amyloid-cancer connection [66].

In light of the loss of ability to form amyloid-like structures by PNT3^3A^, it is conceivable that the lower propensity of NiV V to undergo a liquid-hydrogel transition may reside in the presence of only two contiguous Tyr within the PNT3 region of NiV V instead of three as in HeV V. Whether this has a functional relevance in terms of pathogenicity, immune response evasion and onset and progression of virus-induced encephalopathy remains to be established.

The relative fragility of PNT3 fibers, pinpointed by their sensitivity to SDS, might reflect a role as regulatory switches, i.e. fibers can form and disassemble in response to changes in the surrounding environment and this can play a role in the (in)activation of specific cellular pathways [25, 63, 67]. If fibers correspond to the solid-like structure within droplets resulting from phase separation, then their ability to disassemble might impart metastability (i.e. reversibility) to membrane-less organelles containing such amyloidogenic proteins. Given the link between material properties and pathological conditions [44, 61], cells have likely evolved mechanisms to monitor and control the fluidity of phase-separated droplets. Consistent with this, many ribonucleoprotein (RNP) bodies and granules are enriched in ATP-dependent chaperones such as hsp70 and hsp40 [68]. In agreement, the viscoelasticity of nucleoli and the dynamics of stress granule components exhibit a strong ATP dependence [69, 70]. Our results showing that hsp70 delays/impairs fibril formation by PNT3 are in line with these previous findings, and may reflect a mechanism whereby variations in hsp70 intracellular levels, typically occurring during viral infections (see [71] and references therein cited), may control the stability of phase separated droplets containing the P/V protein. In addition, the ability of hsp70 to hamper fibril formation by PNT3 may also be linked to the well-documented protective role of this chaperone against (paramyxo)viral infections [71-75].

Future efforts will focus on deciphering the underlying molecular mechanisms by which PNT3 fibrils reduces cell resistance to a stress agent in mammal cells. In particular, and in light of recent findings that showed that the Ebola virus VP35 protein (the functional counterpart of paramyxoviral P) blocks stress granule assembly, we will investigate possible interferences with stress granule (dis)assembly. Concomitantly, we will assess any possible synergistic contributions to the phenomenon of phase separation of PNT3 brought either by the C-terminal domain of the V and W proteins, or by oligomerization (i.e. in the context of the P protein).

Since the P protein is an essential component of the replicative complex of *Mononegavirales*, and since the presence of large IDRs is a wide spread and conserved property in *Mononegavirales* P proteins [16, 76-89], it is conceivable that the ability to undergo phase separation and transition is not unique to HeV P/V. As such, these results promise to have broad implications on a large number of important human pathogens.

## Materials and Methods

### Generation of the constructs

The constructs encoding *Henipavirus* V proteins, as well as their Zinc-finger and PNT domains, have already been described [8, 23].

The HeV PNT1, PNT2, PNT3 and PNT4 DNA fragments, encoding respectively aa 1-110, 100-210, 200-310 and 300-404 of the HeV P/V protein, were PCR-amplified using the the pDEST14/HeV PNT construct as template [8] and primers: PNT1-AttB1 and PNT1-AttB2; PNT2-AttB1 and PNT2-AttB2; PNT3-AttB1 and PNT3-AttB2; PNT4-AttB1 and PNT4-AttB2.

The amplicons were then cloned into the pDest17 bacterial expression vector using the Gateway ^®^ technology (Invitrogen). This vector allows expression of recombinant proteins under the control of the T7 promoter. The resulting protein is preceded by a stretch of 22 vector-encoded residues (MSYYHHHHHHLESTSLYKKAGS) encompassing a hexahistidine tag.

For the prokaryotic expression of PNT3 fused the green fluorescent protein (GFP), a 6His-tagged PNT3-GFP construct was generated by PCR amplification using pDest17/PNT3 as template and primers B1HisNT3 and NT3B2. After DpnI treatment, 1 μl of the first PCR was used as a template in a second PCR amplification using primers attl1a and attl2a as described in [90]. The second PCR product was then used in LR reaction with the expression vector pTH31 [91] to yield a construct referred to as His-PNT3-pTH31.

For the eukaryotic expression of PNT3-GFP, another 6His-tagged PNT3-GFP construct was generated by PCR using a GFP coding sequence optimized for eukaryotic expression. In a first PCR, the coding sequence of PNT3 was amplified using His-PNT3-pTH31 as template and primers HindNT3 and NT3R. In a second PCR, the coding sequence of GFP was PCR amplified using GFPq in pTTo [92] as template and primers GFPqF and GFPqXhoI. After DpnI treatment, 1 μl of PCR1 and 1 μl of PCR2 were used as overlapping megaprimers along with primers HindNT3 and GFPqXhoI in a third PCR. After purification, the third PCR product was digested by HindIII and XhoI and ligated into pCDNA3.1+ that had been digested with the same enzymes. The resulting construct is referred to as His-PNT3-GFPq-pCDNA3.1+.

For the eukaryotic expression of PNT3, a 6His-PNT3 construct was generated by PCR using His-PNT3-GFPq-pCDNA3.1+ as template and primers HindNT3 and NT3XhoI. After DpnI treatment, the PCR product was digested by HindIII and XhoI and ligated to pCDNA3.1+ as described above. The construct is referred to as His-PNT3-pCDNA3.1+.

For the prokaryotic expression of the PNT3 Y210A-Y211A-Y212A triple variant (PNT3^3A^), the His-PNT3-pTH31 construct was used as template in two separate PCR amplifications using either primers attB1 and R_3ala-pNT3 (PCR1), or primers F_3ala-pNT3 and attB2 (PCR2). After DpnI treatment, 1 μl of PCR1 and 1 μl of PCR2 were used as overlapping megaprimers along with primers attB1 and attB2 in a third PCR. After purification, the third PCR product was inserted in pDONR by BP reaction (Invitrogen). After sequencing, the mutant coding sequence was transferred from pDONR to pDEST17 by LR reaction (Invitrogen). The resulting construct is referred to as mutPNT3-pDEST17.

For the eukaryotic expression of the PNT3 Y210A-Y211A-Y212A triple variant, the His-PNT3-pCDNA3.1+ construct was used as template in two separate PCR amplifications using either primers HindNT3 and R_3ala-pNT3 (PCR1), or primers F_3ala-pNT3 and NT3XhoI (PCR2). After DpnI treatment, 1 μl of PCR1 and 1 μl of PCR2 were used as overlapping megaprimers along with primers HindNT3 and NT3XhoI in a third PCR. After purification, the third PCR product was processed as described for generating His-PNT3-3A-pCDNA3.1+.

The constructs for the eukaryotic expression of GFP and 2OST (an irrelevant, control hexahistidine tagged protein) contain the ORF of GFPq [92] and of 2OST, respectively, cloned into pCDNA3.1+. The construct used for the prokaryotic expression of His-hsp70 has already been described [93].

Primers were all purchased from Eurofins Genomics. The sequence of all the primers is provided in **Supplementary Table S2**. All the constructs were verified by DNA sequencing (Eurofins Genomics) and found to conform to expectations.

### Proteins expression and purification

The *E. coli* strain T7pRos was used for the expression of all the recombinant proteins. Cultures were grown over-night to saturation in LB medium containing 100 μg mL^−1^ ampicillin and 34 μg mL^−1^ chloramphenicol. An aliquot of the overnight culture was diluted 1/20 into 1 L of TB medium and grown at 37 °C 200 rpm. When the optical density at 600 nm (OD_600_) reached 0.5-0.8, isopropyl β-D-thiogalactopyanoside (IPTG) was added to a final concentration of 0.5 mM, and the cells were grown at 25 °C 200 rpm overnight. The induced cells were harvested, washed and collected by centrifugation (5,000 rpm, 10 min). The resulting pellets were frozen at −20 °C.

Expression of ^15^N-labelled PNT3 was performed as described above except that when the cultures reached an OD_600_ of 0.6, the culture was centrifuged at 4,000 rpm for 10 min, and the pellet was resuspended in 250 mL of M9 medium (6 g L^−1^ of Na_2_HPO_4_, 3 g L^−1^ of KH_2_PO_4_, 0.5 g L^−1^ of NaCl, 0.246 g L^−1^ of MgSO_4_) supplemented with 1 g L^−1^ of ^15^NH_4_Cl and 4 g L^−1^ of glucose. After one hour at 37 °C, IPTG was added to a final concentration of 0.5 mM, and the cells were subsequently grown at 25 °C overnight.

The purification protocol of the *Henipavirus* V proteins and of the HeV V ZnFD have been already reported [23], as was that of HeV PNT [8]. The PNT1-PNT4, PNT3^3A^ and PNT3-GFP proteins were purified as follows. The cellular pellet was resuspended (30 mL *per* liter of culture) in buffer A (50 mM Tris/HCl pH 7.5, 500 mM NaCl, 20 mM imidazole) supplemented with 1 mM phenyl methyl sulfonyl fluoride (PMSF), 0.1 mg mL^−1^ lysozyme, 10 μg mL^−1^ DNAse I and 20 mM MgSO_4_. After an incubation of 20 min at 4 °C, the cells were disrupted by sonication (using a 750 W sonicator and 3 cycles of 30 s each at 45 % power output). Except for the PNT3-GFP protein, 6 M GuHCl was added to the lysate and the solution was incubated 1 hour at 4 °C with gentle agitation. The lysates were clarified by centrifugation at 14,000 g for 30 min at 4 °C and the supernatant was loaded onto a 5 mL Nickel column (GE Healthcare) pre-equilibrated in buffer A. The affinity resin was washed with 20 column volumes (CV) of buffer A. Proteins were eluted with buffer A supplemented with 250 mM imidazole. PNT3 and PNT3^3A^ were concentrated in the presence of 6 M GuHCl up to 1-2 mM using Centricon concentrators (Merk-Millipore) with a 10 kDa molecular mass cut-off, and then frozen at −20 °C.

Prior to each experiment, the proteins were subjected to size exclusion chromatography (SEC) that also enabled confirming homogeneity and monomeric nature of the proteins. The SEC column was equilibrated with buffer B (20 mM Tris/HCl pH 7.5, 100 mM NaCl, 5 mM EDTA). For samples to be used in CD, SAXS, EM and NMR experiments, SEC was carried out using a 50 mM phosphate pH 6.5, 5 mM EDTA buffer.

Fractions from all purification steps were analyzed by SDS-PAGE. Protein concentrations were estimated using the theoretical absorption coefficients at 280 nm as obtained using the program ProtParam from the EXPASY server.

The purity of the final purified products was assessed by SDS-PAGE (**Figure 1D**). The identity of the purified PNT3 and PNT3^3A^ samples was confirmed by mass spectrometry (MALDI-TOF) analysis of tryptic fragments obtained after digestion of the purified protein bands excised from SDS-polyacrylamide gels (**Supplementary Figure S4**). The bands were then digested with trypsin (20 μl of trypsin/band at a concentration of 12.5 ng/μl). For each protein band, mass analyses were performed on a MALDI-TOF-TOF Bruker Ultraflex III spectrometer (Bruker Daltonics, Wissembourg, France) operating in reflector mode. The *m/z* range was from 600 to 4500, and α -Cyano-4-hydroxycinnamic acid was used as matrix solution. The mass standards were either autolytic tryptic peptides used as internal standards or peptide standards (Bruker Daltonics). Following MS analysis, MS/MS analyses were performed on the most intense peaks to identify the amino acid sequence of the protein band.

Hsp70 was purified according to [94] except that the last SEC step was replaced by a desalting step using a HiPrep 26/10 desalting column (GE Healthcare) and buffer B. The protein was then concentrated to 0.5 mM using Centricon concentrators (Merk-Millipore) with a 30 kDa molecular mass cut-off.

### Turbidity and aggregation measurements

Turbidity was measured by monitoring the OD at 600 nm using a NanoDrop ND-1000 (Thermo Scientific) spectrophotometer. Samples (100 μL each), in the concentration range of 0-250 μM, were incubated 1-hour at RT either in the absence or in the presence of increasig concentrations of the crowding agent PEG300. Different PEG (400, 4000, 6000, 8000) at different percentages (3-30 %) were also tested and the evolution of the droplets (400 nL) were monitored using a visible/UV light RockImager (Formulatrix). The formation of aggregates was monitored by measuring the OD at 340 nm of PNT3 samples incubated 1 hour either at RT or at 370C in the presence of increasing concentrations of salt.

### Thioflavin T and Congo red binding assays

Thioflavin T (ThT) (Sigma-Aldrich) binding assays were performed using the dye at a final concentration of 20 μM in the presence of sodium azide 0.05 %. The samples were excited at 440 nm and emission was recorded between 450 and 500 nm. Fluorescence intensity was measured over time at each 15 min at 37 °C using a Cary Eclipse Fluorescence Spectrophotometer (Agilent). Data was plotted using Prism software and were analyzed using the Amylofit server (www.amylofit.ch.cam.ac.uk) [95].

CR binding assays were performed using the dye at a final concentration of 5 μM in the presence of 25 μM PNT3 (in sodium phosphate buffer at pH 6.5). The spectrum was recorded after 27 hours of incubation at 37°C using a NanoDrop ND-1000 (Thermo Scientific) spectrophotometer in the 400–700 nm range. Five μM CR in sodium phosphate buffer without PNT3 was used as a control.

### Far-UV circular dichroism

CD spectra were measured using a Jasco 819 dichrograph, flushed with N_2_ and equipped with a Peltier thermoregulation system. Proteins were loaded into a 1 mm quartz cuvette at 0.1 mg/mL (in 10 mM phosphate buffer at pH 6.5) and spectra were recorded at 20 °C. The scanning speed was 20 nm min^−1^, with data pitch of 0.2 nm. Each spectrum is the average of three acquisitions. The spectrum of buffer was subtracted from the protein spectrum. Spectra were smoothed using the “means-movement” smoothing procedure implemented in the Spectra Manager package.

Mean molar ellipticity values per residue (MRE) were calculated as

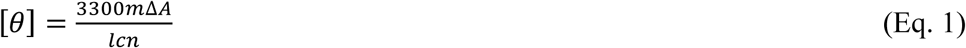

where *l* is the path length in cm, *n* is the number of residues (133), *m* is the molecular mass in Daltons (15,138) and *c* is the concentration of the protein in mg mL^−1^.

### Small-angle X-ray scattering (SAXS)

SAXS measurements were carried out at the European Synchrotron Radiation Facility (Grenoble, France) on beamline BM29 (bending magnet) at a working energy of 12.5 KeV. Data were collected on a Pilatus (1M) detector. The wavelength was 0.992 Å. The sample-to-detector distance was 2.847 m, leading to scattering vectors (q) ranging from 0.028 to 4.525 nm^−1^. The scattering vector is defined as q = 4π/λsinθ, where 2θ is the scattering angle. The exposure time was optimized to reduce radiation damage. SAXS data were collected at 20 °C using purified protein samples (50 μL each). PNT3 concentrations were 1.0 and 2.0 mg mL^−1^. Samples were incubated at 3°C and the data were collected at various times points from 0 to 630 min. Data were analyzed using the ATSAS program package [96]. Data reductions were performed using the established procedure available at BM29, and buffer background runs were subtracted from sample runs. The forward scattering intensities were calibrated using water as reference.

The R_g_, forward intensity at zero angle I(0) and pair distance distribution function, P(r), were calculated with the program GNOM [97]. This enabled identifying the scattering curves corresponding to a monomeric species. For those latter curves, the R_g_ and forward intensity at zero angle I(0) were recalculated and refined with the program PRIMUS [98] according to the Guinier approximation at low q values, in a q.Rg range up to 1.3:

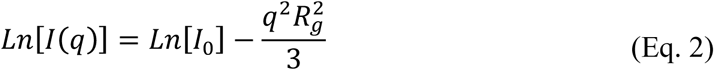

For the monomeric species at the highest protein concentration, we also attempted at describing it as conformational ensemble. To this end we used the program suite EOM 2.0 [42] using the default parameters.

The theoretical value of Rg (in Å) expected for an IDP was calculated using Flory’s equation according to [41]:

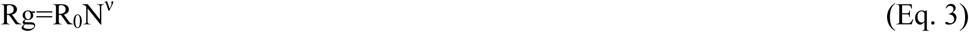

where N is the number of amino acid residues, R_0_ is 2.54 ± 0.01 and v is 0.522 ± 0.01

### Negative-staining transmission electron microscopy (TEM)

Drops of 2 μL of freshly purified PNT3 or PNT^3A^ proteins (100-200 μM) either in the absence or in the presence of a two-fold molar excess of Hsp70, were deposited onto a glow-discharge carbon coated grid (Formwar/Carbon 300 mesh Cu, Agar Scientific). Prior to protein deposition, grids were exposed to plasma glow discharge for 20 seconds using a PELCO, easiGlow Glow Discharge Cleaning System (Ted Pella Inc. USA) (Current 25mA) in order to increase protein adhesion. To assess the fibrils stability, PNT3 (200 μM) supplemented with 5 mM SDS was deposited on the copper grids. The grids were washed three times with 20 μL of deionized water before incubating them for 1 min in a 2 % (w/v) Uranyl Acetate solution (LauryLab-France). Images were collected using a TECNAI T12 Spirit microscope (FEI company) operated at 120 kV and an Eagle 2Kx2K CCD camera (FEI company).

### Nuclear Magnetic Resonance (NMR)

A sample of a freshly purified ^15^N-labelled PNT3 at 0.1 mM in 50 mM phosphate pH 6.5, 5 mM EDTA, also containing D2O (10 %), was used for the acquisition of 1D ^1^H and 2D ^1^H,^15^N HSQC spectra with an 22.3 T Bruker AvanceIII 950 ultra-shielded-plus spectrometer equipped with a triple resonance CryoProbe (TCI) at 310 K. The sample was incubated at 37°C and spectra were recorded at various time points. Spectra were recorded both at 288 and 310 K and were found to be superimposable.

### X-ray diffraction

A sample of PNT3 at 20 μM from a ThT assay was centrifuged at 13000 rpm for 15 min and lyophilized. The powder was transferred into a mounted cryo-loop (Hampton Research). The fiber diffraction images were collected using a Microstar X-ray generator equipped with a MAR345 detector using a fixed wavelength of 1.54 Å and a sample-detector distance of 200 mm, with an exposure time of 300 s.

### Assays of SDS Sensitivity

Preformed fibers of PNT3 (100 μM after 56 hours of incubation at 37°C) either non-supplemented or supplemented with 2% SDS, were passed through a 0.2 mm spin filter to remove fiber particles. UV absorbance, after subtraction of buffer contributions, was measured to monitor the amount of monomeric protein that passed through the filter.

### Transfection of mammalian cells and viability assays

HEK293T cells were cultured in DMEM supplemented with 10% FBS at 37°C in a CO_2_ incubator. Transfections were performed with PEI reagent (Polysciences) with a DNA:PEI ratio of 1:2 (w/w). CR staining and viability assay experiments were performed 72 and 48 hours after transfection, respectively. For CR staining, cells were fixed with 4 % PFA and incubated over night with 5 μM CR (Sigma) in a buffer containing 100 mM NaCl, 10 mM Hepes pH 7.4. Cells were then washed 1 hour with PBS diluted in water (1/4) and stored at 4°C in PBS before microscopy analysis. For cell viability assays, non-transfected or transfected (6His-PNT3, 6His-PNT3^3A^, 6His-PNT3-GFP, GFP and 6His-2OST) cells were incubated 6 hours with H2O2 at different concentrations in triplicate in 96-well plates. PrestoBlue reagent (ThermoFisher) was added (10% v/v) to the 96-well plate and then incubated 2 additional hours at 37°C. Fluorescence was measured (ex 560 nm / em 590 nm) with a Saphire2 fluorimeter (Tecan). Cells expressing GFP or 6His-PNT3-GFP were observed under an epi-fluorescent microscope (Eclipse TS-100 Nikon).

### Western Blot analysis

Expression 6His-PNT3, 6His-PNT33A, and 6His-2OST in HEK293T cells was assessed 48 hours post-transfection in 96-well plates. Cells were scrapped off after trypsinization (96 wells) and 5X Laemli loading buffer was added. Samples were boiled and 20 *μ* L of lysates were loaded onto a 15% SDS-PAGE gel. After electrophoresis, the gel was transferred on a nitrocellulose membrane and the proteins were stained using a Penta-His HRP conjugate (1/2000) (Biolegend). Signal was detected by chemiluminescence (Amersham ECL Western Blotting detecting reagent, GE Healthcare) using a Kodak, Image Station 440 luminometer.

## Supporting information

## Acknowledgements

This work was carried out with the financial support of the CNRS. It was also partly supported by the French Infrastructure for Integrated Structural Biology (FRISBI) (ANR-10-INSB-05-01). E.S. is supported by a joint doctoral fellowship from the Direction Générale de l'Armement (DGA) and Aix-Marseille University.

We thank Patrick Fourquet, from the mass spectrometry platform of the Centre de Recherche en Cancérologie de Marseille (CRCM) for mass spectrometry analyses and Petra Pernot for her help in SAXS data collection, and the ESRF synchrotron for beamtime allocation. Finally, we want to express our gratitude to Philippe Cantau (AFMB lab) for useful assistance with X-ray diffraction and Adéline Goulet (AFMB lab) for useful discussions on negative-staining electron microscopy. We are also grateful to Gerlind Sulzenbacher (AFMB lab) for efficiently managing the AFMB BAG. We thank Frédéric Carrière (BIP lab, Marseille) for useful comments on PNT3 phase separation. We thank Dr Helena Berglund for kindly providing us with pTH31.

## References

[1] Wang LF, Yu M, Hansson E, Pritchard LI, Shiell B, Michalski WP, et al. The exceptionally large genome of Hendra virus: support for creation of a new genus within the family Paramyxoviridae. J Virol. 2000;74:9972–9.

[2] Gurley ES, Montgomery JM, Hossain MJ, Bell M, Azad AK, Islam MR, et al. Person-to-person transmission of Nipah virus in a Bangladeshi community. Emerg Infect Dis. 2007;13:1031–7.

[3] Homaira N, Rahman M, Hossain MJ, Epstein JH, Sultana R, Khan MS, et al. Nipah virus outbreak with person-to-person transmission in a district of Bangladesh, 2007. Epidemiol Infect. 2010;138:1630–6.

[4] Ching PK, de los Reyes VC, Sucaldito MN, Tayag E, Columna-Vingno AB, Malbas FF, Jr., et al. Outbreak of henipavirus infection, Philippines, 2014. Emerg Infect Dis. 2015;21:328–31.

[5] Hayman DT, Suu-Ire R, Breed AC, McEachern JA, Wang L, Wood JL, et al. Evidence of henipavirus infection in West African fruit bats. PLoS One. 2008;3:e2739.

[6] Bloyet LM, Welsch J, Enchery F, Mathieu C, de Breyne S, Horvat B, et al. HSP90 Chaperoning in Addition to Phosphoprotein Required for Folding but Not for Supporting Enzymatic Activities of Measles and Nipah Virus L Polymerases. J Virol. 2016;90:6642–56.

[7] Sourimant J, Rameix-Welti MA, Gaillard AL, Chevret D, Galloux M, Gault E, et al. Fine mapping and characterization of the L-polymerase-binding domain of the respiratory syncytial virus phosphoprotein. J Virol. 2015;89:4421–33.

[8] Habchi J, Mamelli L, Darbon H, Longhi S. Structural Disorder within Henipavirus Nucleoprotein and Phosphoprotein: From Predictions to Experimental Assessment. PLoS ONE. 2010;5:e11684.

[9] Blocquel D, Beltrandi M, Erales J, Barbier P, Longhi S. Biochemical and structural studies of the oligomerization domain of the Nipah virus phosphoprotein: Evidence for an elongated coiled-coil homotrimer. Virology. 2013;446:162–72.

[10] Beltrandi M, Blocquel D, Erales J, Barbier P, Cavalli A, Longhi S. Insights into the coiled-coil organization of the Hendra virus phosphoprotein from combined biochemical and SAXS studies. Virology. 2015;477:42–55.

[11] Bruhn-Johannsen JF, Barnett K, Bibby J, Thomas J, Keegan R, Rigden D, et al. Crystal structure of the Nipah virus phosphoprotein tetramerization domain. J Virol. 2014;88:758–62.

[12] Communie G, Habchi J, Yabukarski F, Blocquel D, Schneider R, Tarbouriech N, et al. Atomic resolution description of the interaction between the nucleoprotein and phosphoprotein of Hendra virus. PLoS Pathog. 2013;9:e1003631.

[13] Habchi J, Blangy S, Mamelli L, Ringkjobing Jensen M, Blackledge M, Darbon H, et al. Characterization of the interactions between the nucleoprotein and the phosphoprotein of Henipaviruses. J Biol Chem. 2011;286:13583–602.

[14] Habchi J, Martinho M, Gruet A, Guigliarelli B, Longhi S, Belle V. Monitoring structural transitions in IDPs by site-directed spin labeling EPR spectroscopy. Methods Mol Biol. 2012;895:361–86.

[15] Blocquel D, Habchi J, Gruet A, Blangy S, Longhi S. Compaction and binding properties of the intrinsically disordered C-terminal domain of Henipavirus nucleoprotein as unveiled by deletion studies. Mol Biosyst. 2012;8:392–410.

[16] Karlin D, Ferron F, Canard B, Longhi S. Structural disorder and modular organization in Paramyxovirinae N and P. J Gen Virol. 2003;84:3239–52.

[17] Fontana JM, Bankamp B, Rota PA. Inhibition of interferon induction and signaling by paramyxoviruses. Immunol Rev. 2008;225:46–67.

[18] Audsley MD, Moseley GW. Paramyxovirus evasion of innate immunity: Diverse strategies for common targets. World J Virol. 2013;2:57–70.

[19] Park MS, Shaw ML, Munoz-Jordan J, Cros JF, Nakaya T, Bouvier N, et al. Newcastle disease virus (NDV)-based assay demonstrates interferon-antagonist activity for the NDV V protein and the Nipah virus V, W, and C proteins. J Virol. 2003;77:1501–11.

[20] Shaw ML, Garcia-Sastre A, Palese P, Basler CF. Nipah virus V and W proteins have a common STAT1-binding domain yet inhibit STAT1 activation from the cytoplasmic and nuclear compartments, respectively. J Virol. 2004;78:5633–41.

[21] Marsh GA, de Jong C, Barr JA, Tachedjian M, Smith C, Middleton D, et al. Cedar virus: a novel Henipavirus isolated from Australian bats. PLoS Pathog. 2012;8:e1002836.

[22] Ulane CM, Horvath CM. Paramyxoviruses SV5 and HPIV2 assemble STAT protein ubiquitin ligase complexes from cellular components. Virology. 2002;304:160–6.

[23] Salladini E, Delauzun V, Longhi S. The Henipavirus V protein is a prevalently unfolded protein with a zinc-finger domain involved in binding to DDB1. Mol Biosyst. 2017;13:2254–67.

[24] Uversky VN. Intrinsically disordered proteins in overcrowded milieu: Membrane-less organelles, phase separation, and intrinsic disorder. Curr Opin Struct Biol. 2017;44:18–30.

[25] Holehouse AS, Pappu RV. Functional Implications of Intracellular Phase Transitions. Biochemistry. 2018;57:2415–23.

[26] Boeynaems S, Alberti S, Fawzi NL, Mittag T, Polymenidou M, Rousseau F, et al. Protein Phase Separation: A New Phase in Cell Biology. Trends Cell Biol. 2018.

[27] Murray DT, Kato M, Lin Y, Thurber KR, Hung I, McKnight SL, et al. Structure of FUS Protein Fibrils and Its Relevance to Self-Assembly and Phase Separation of Low-Complexity Domains. Cell. 2017;171:615–27 e16.

[28] Kato M, McKnight SL. A Solid-State Conceptualization of Information Transfer from Gene to Message to Protein. Annu Rev Biochem. 2017.

[29] Boeynaems S, Bogaert E, Kovacs D, Konijnenberg A, Timmerman E, Volkov A, et al. Phase Separation of C9orf72 Dipeptide Repeats Perturbs Stress Granule Dynamics. Mol Cell. 2017;65:1044–55 e5.

[30] Maharana S, Wang J, Papadopoulos DK, Richter D, Pozniakovsky A, Poser I, et al. RNA buffers the phase separation behavior of prion-like RNA binding proteins. Science. 2018.

[31] Lin Y, Currie SL, Rosen MK. Intrinsically disordered sequences enable modulation of protein phase separation through distributed tyrosine motifs. J Biol Chem. 2017;292:19110–20.

[32] Vernon RM, Chong PA, Tsang B, Kim TH, Bah A, Farber P, et al. Pi-Pi contacts are an overlooked protein feature relevant to phase separation. Elife. 2018;7.

[33] Li HR, Chiang WC, Chou PC, Wang WJ, Huang JR. TAR DNA-binding protein 43 (TDP-43) liquid-liquid phase separation is mediated by just a few aromatic residues. J Biol Chem. 2018;293:6090–8.

[34] Galzitskaya OV. Repeats are one of the main characteristics of RNA-binding proteins with prion-like domains. Mol Biosyst. 2015;11:2210–8.

[35] Axelrod D, Koppel DE, Schlessinger J, Elson E, Webb WW. Mobility measurement by analysis of fluorescence photobleaching recovery kinetics. Biophys J. 1976;16:1055–69.

[36] Klunk WE, Pettegrew JW, Abraham DJ. Quantitative evaluation of congo red binding to amyloid-like proteins with a beta-pleated sheet conformation. J Histochem Cytochem. 1989;37:1273–81.

[37] LeVine H, 3rd. Thioflavine T interaction with synthetic Alzheimer’s disease beta-amyloid peptides: detection of amyloid aggregation in solution. Protein Sci. 1993;2:404–10.

[38] Sunde M, Blake C. The structure of amyloid fibrils by electron microscopy and X-ray diffraction. Adv Protein Chem. 1997;50:123–59.

[39] Stromer T, Serpell LC. Structure and morphology of the Alzheimer’s amyloid fibril. Microsc Res Tech. 2005;67:210–7.

[40] Morris KL, Serpell LC. X-ray fibre diffraction studies of amyloid fibrils. Methods Mol Biol. 2012;849:121–35.

[41] Bernado P, Blackledge M. A self-consistent description of the conformational behavior of chemically denatured proteins from NMR and small angle scattering. Biophys J. 2009;97:2839–45.

[42] Tria G, Mertens HDT, Kachala M, Svergun D. Advanced ensemble modelling of flexible macromolecules using X-ray solution scattering. IUCrJ. 2015;2:202–17.

[43] Dobson CM. Protein folding and misfolding. Nature. 2003;426:884–90.

[44] Chiti F, Dobson CM. Protein Misfolding, Amyloid Formation, and Human Disease: A Summary of Progress Over the Last Decade. Annu Rev Biochem. 2017;86:27–68.

[45] Alberti S, Halfmann R, King O, Kapila A, Lindquist S. A systematic survey identifies prions and illuminates sequence features of prionogenic proteins. Cell. 2009;137:146–58.

[46] Kato M, Han TW, Xie S, Shi K, Du X, Wu LC, et al. Cell-free formation of RNA granules: low complexity sequence domains form dynamic fibers within hydrogels. Cell. 2012;149:753–67.

[47] Fenyi A, Coens A, Bellande T, Melki R, Bousset L. Assessment of the efficacy of different procedures that remove and disassemble alpha-synuclein, tau and A-beta fibrils from laboratory material and surfaces. Sci Rep. 2018;8:10788.

[48] Klucken J, Shin Y, Masliah E, Hyman BT, McLean PJ. Hsp70 Reduces alpha-Synuclein Aggregation and Toxicity. J Biol Chem. 2004;279:25497–502.

[49] Dedmon MM, Christodoulou J, Wilson MR, Dobson CM. Heat shock protein 70 inhibits alpha-synuclein fibril formation via preferential binding to prefibrillar species. J Biol Chem. 2005;280:14733–40.

[50] Patterson KR, Ward SM, Combs B, Voss K, Kanaan NM, Morfini G, et al. Heat shock protein 70 prevents both tau aggregation and the inhibitory effects of preexisting tau aggregates on fast axonal transport. Biochemistry. 2011;50:10300–10.

[51] Aprile FA, Arosio P, Fusco G, Chen SW, Kumita JR, Dhulesia A, et al. Inhibition of alpha-Synuclein Fibril Elongation by Hsp70 Is Governed by a Kinetic Binding Competition between alpha-Synuclein Species. Biochemistry. 2017;56:1177–80.

[52] Xue YL, Wang H, Riedy M, Roberts BL, Sun Y, Song YB, et al. Molecular dynamics simulations of Hsp40 J-domain mutants identifies disruption of the critical HPD-motif as the key factor for impaired curing in vivo of the yeast prion [URE3]. J Biomol Struct Dyn. 2018;36:1764–75.

[53] Lahaye X, Vidy A, Pomier C, Obiang L, Harper F, Gaudin Y, et al. Functional characterization of Negri bodies (NBs) in rabies virus-infected cells: Evidence that NBs are sites of viral transcription and replication. J Virol. 2009;83:7948–58.

[54] Heinrich BS, Cureton DK, Rahmeh AA, Whelan SP. Protein expression redirects vesicular stomatitis virus RNA synthesis to cytoplasmic inclusions. PLoS Pathog. 2010;6:e1000958.

[55] Hoenen T, Shabman RS, Groseth A, Herwig A, Weber M, Schudt G, et al. Inclusion bodies are a site of ebolavirus replication. J Virol. 2012;86:11779–88.

[56] Zhang S, Chen L, Zhang G, Yan Q, Yang X, Ding B, et al. An amino acid of human parainfluenza virus type 3 nucleoprotein is critical for template function and cytoplasmic inclusion body formation. J Virol. 2013;87:12457–70.

[57] Rincheval V, Lelek M, Gault E, Bouillier C, Sitterlin D, Blouquit-Laye S, et al. Functional organization of cytoplasmic inclusion bodies in cells infected by respiratory syncytial virus. Nat Commun. 2017;8:563.

[58] Cifuentes-Munoz N, Branttie J, Slaughter KB, Dutch RE. Human Metapneumovirus Induces Formation of Inclusion Bodies for Efficient Genome Replication and Transcription. J Virol. 2017;91.

[59] Netherton CL, Wileman T. Virus factories, double membrane vesicles and viroplasm generated in animal cells. Curr Opin Virol. 2011;1:381–7.

[60] Nikolic J, Le Bars R, Lama Z, Scrima N, Lagaudriere-Gesbert C, Gaudin Y, et al. Negri bodies are viral factories with properties of liquid organelles. Nat Commun. 2017;8:58.

[61] Murakami T, Qamar S, Lin JQ, Schierle GS, Rees E, Miyashita A, et al. ALS/FTD Mutation-Induced Phase Transition of FUS Liquid Droplets and Reversible Hydrogels into Irreversible Hydrogels Impairs RNP Granule Function. Neuron. 2015;88:678–90.

[62] Patel A, Lee HO, Jawerth L, Maharana S, Jahnel M, Hein MY, et al. A Liquid-to-Solid Phase Transition of the ALS Protein FUS Accelerated by Disease Mutation. Cell. 2015;162:1066–77.

[63] Elbaum-Garfinkle S, Brangwynne CP. Liquids, Fibers, and Gels: The Many Phases of Neurodegeneration. Dev Cell. 2015;35:531–2.

[64] Alonso LG, Garcia-Alai MM, Smal C, Centeno JM, Iacono R, Castano E, et al. The HPV16 E7 viral oncoprotein self-assembles into defined spherical oligomers. Biochemistry. 2004;43:3310–7.

[65] Smal C, Alonso LG, Wetzler DE, Heer A, de Prat Gay G. Ordered self-assembly mechanism of a spherical oncoprotein oligomer triggered by zinc removal and stabilized by an intrinsically disordered domain. PLoS One. 2012;7:e36457.

[66] Dantur K, Alonso L, Castano E, Morelli L, Centeno-Crowley JM, Vighi S, et al. Cytosolic accumulation of HPV16 E7 oligomers supports different transformation routes for the prototypic viral oncoprotein: the amyloid-cancer connection. Int J Cancer. 2009;125:1902–11.

[67] Shin Y, Brangwynne CP. Liquid phase condensation in cell physiology and disease. Science. 2017;357.

[68] Protter DS, Parker R. Principles and Properties of Stress Granules. Trends Cell Biol. 2016;26:668–79.

[69] Jain S, Wheeler JR, Walters RW, Agrawal A, Barsic A, Parker R. ATPase-Modulated Stress Granules Contain a Diverse Proteome and Substructure. Cell. 2016;164:487–98.

[70] Brangwynne CP, Mitchison TJ, Hyman AA. Active liquid-like behavior of nucleoli determines their size and shape in Xenopus laevis oocytes. Proc Natl Acad Sci U S A. 2011;108:4334–9.

[71] Oglesbee M, Kim MY, Shu Y, Longhi S. Extracellular HSP70, Neuroinflammation and Protection against Viral Virulence. In: Asea AAAaK, P, editor. Chaperokine activity of heat shock proteins: Springer; 2018.

[72] Kim MY, Shu Y, Carsillo T, Zhang J, Yu L, Peterson C, et al. hsp70 and a novel axis of type I interferon-dependent antiviral immunity in the measles virus-infected brain. J Virol. 2013;87:998–1009.

[73] Oglesbee MJ, LaBranche T. Inside-out: extracellular roles for heat shock proteins. Vet Pathol. 2013;50:921–4.

[74] Kim MY, Ma Y, Zhang Y, Li J, Shu Y, Oglesbee M. hsp70-dependent antiviral immunity against cytopathic neuronal infection by vesicular stomatitis virus. J Virol. 2013;87:10668–78.

[75] Ma Y, Duan Y, Wei Y, Liang X, Niewiesk S, Oglesbee M, et al. Heat shock protein 70 enhances mucosal immunity against human norovirus when coexpressed from a vesicular stomatitis virus vector. J Virol. 2014;88:5122–37.

[76] Bourhis JM, Canard B, Longhi S. Structural disorder within the replicative complex of measles virus: functional implications. Virology. 2006;344:94–110.

[77] Longhi S, Oglesbee M. Structural disorder within the measles virus nucleoprotein and phosphoprotein. Protein and Peptide Letters. 2010;17:961–78.

[78] Leyrat C, Gerard FC, de Almeida Ribeiro E, Jr., Ivanov I, Ruigrok RW, Jamin M. Structural disorder in proteins of the rhabdoviridae replication complex. Protein Pept Lett. 2010;17:979–87.

[79] Longhi S. Structural disorder within the measles virus nucleoprotein and phosphoprotein: functional implications for transcription and replication. In: Luo M, editor. Negative strand RNA virus. Singapore: World Scientific Publishing; 2011. p. 95–125.

[80] Ivanov I, Yabukarski F, Ruigrok RW, Jamin M. Structural insights into the rhabdovirus transcription/replication complex. Virus Res. 2011;162:126–37.

[81] Leyrat C, Schneider R, Ribeiro EA, Jr., Yabukarski F, Yao M, Gerard FC, et al. Ensemble structure of the modular and flexible full-length vesicular stomatitis virus phosphoprotein. J Mol Biol. 2012;423:182–97.

[82] Blocquel D, Bourhis JM, Eléouët JF, Gerlier D, Habchi J, Jamin M, et al. Transcription et réplication des Mononégavirales: une machine moléculaire originale. Virologie. 2012;16:225–57.

[83] Habchi J, Mamelli L, Longhi S. Structural disorder within the nucleoprotein and phosphoprotein from measles, Nipah and Hendra viruses. In: Uversky VN, Longhi S, editors. Flexible viruses: structural disorder in viral proteins. Hoboken, New Yersey: John Wiley and Sons; 2012. p. 47–94.

[84] Habchi J, Longhi S. Structural disorder within paramyxovirus nucleoproteins and phosphoproteins. Mol Biosyst. 2012;8:69–81.

[85] Communie G, Ruigrok RW, Jensen MR, Blackledge M. Intrinsically disordered proteins implicated in paramyxoviral replication machinery. Curr Opin Virol. 2014;5:72–81.

[86] Leung DW, Borek D, Luthra P, Binning JM, Anantpadma M, Liu G, et al. An Intrinsically Disordered Peptide from Ebola Virus VP35 Controls Viral RNA Synthesis by Modulating Nucleoprotein-RNA Interactions. Cell Rep. 2015;11:376–89.

[87] Erales J, Blocquel D, Habchi J, Beltrandi M, Gruet A, Dosnon M, et al. Order and disorder in the replicative complex of paramyxoviruses. Adv Exp Med Biol. 2015;870:351–81.

[88] Habchi J, Longhi S. Structural Disorder within Paramyxoviral Nucleoproteins and Phosphoproteins in Their Free and Bound Forms: From Predictions to Experimental Assessment. Int J Mol Sci. 2015;16:15688–726.

[89] Longhi S, Bloyet LM, Gianni S, Gerlier D. How order and disorder within paramyxoviral nucleoproteins and phosphoproteins orchestrate the molecular interplay of transcription and replication. Cell Mol Life Sci. 2017;74:3091–118.

[90] Gruet A, Dosnon M, Blocquel D, Brunel J, Gerlier D, Das RK, et al. Fuzzy regions in an intrinsically disordered protein impair protein-protein interactions. FEBS J. 2016;283:576–94.

[91] van den Berg S, Lofdahl PA, Hard T, Berglund H. Improved solubility of TEV protease by directed evolution. J Biotechnol. 2006;121:291–8.

[92] Durocher Y, Perret S, Kamen A. High-level and high-throughput recombinant protein production by transient transfection of suspension-growing human 293-EBNA1 cells. Nucleic Acids Res. 2002;30:E9.

[93] Zhang X, Bourhis JM, Longhi S, Carsillo T, Buccellato M, Morin B, et al. Hsp72 recognizes a P binding motif in the measles virus N protein C-terminus. Virology. 2005;337:162–74.

[94] Couturier M, Buccellato M, Costanzo S, Bourhis JM, Shu Y, Nicaise M, et al. High Affinity Binding between Hsp70 and the C-Terminal Domain of the Measles Virus Nucleoprotein Requires an Hsp40 Co-Chaperone. J Mol Recognit. 2010;23:301–15.

[95] Meisl G, Kirkegaard JB, Arosio P, Michaels TC, Vendruscolo M, Dobson CM, et al. Molecular mechanisms of protein aggregation from global fitting of kinetic models. Nat Protoc. 2016;11:252–72.

[96] Petoukhov MV, Franke D, Shkumatov AV, Tria G, Kikhney AG, Gajda M, et al. New developments in the ATSAS program package for small-angle scattering data analysis. J Appl Cryst 2012;45:342–50.

[97] Svergun D. Determination of the regularization parameters in indirect-trasform methods using perceptual criteria. J Appl Cryst. 1992;25:495–503.

[98] Konarev PV, Volkov VV, Sokolova AV, Koch MHJ, Svergun DI. PRIMUS: a Windows PC-based system for small-angle scattering data analysis. J Appl Cryst. 2003;36:1277–82.

