## Supplementary material for "Phase transition and amyloid formation by a viral protein as an additional molecular mechanism of virus-induced cell toxicity"

Supplementary Figures S1 to S4
Supplementary Tables S1 and S2

### Supplementary Figures

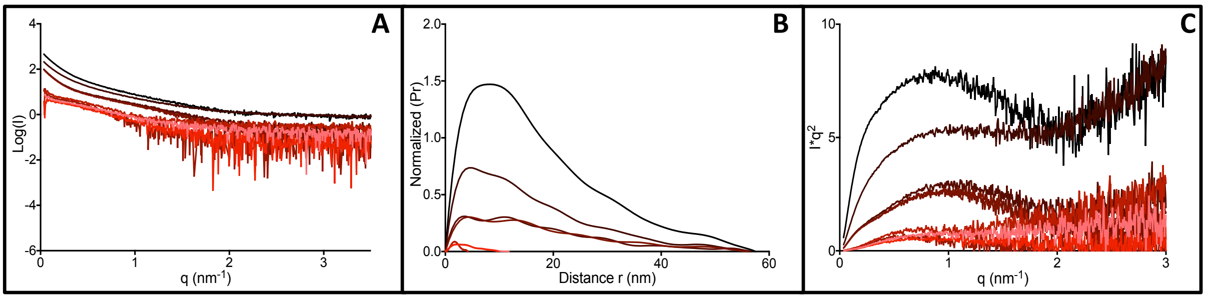


**Supplementary Figure S1. Small-angle X-ray scattering studies of PNT3.** (**A**) Scattering curves as obtained from a sample of PNT3 at 1 mg mL$^{-1}$ at various times of incubation at 37°C. The curves are represented with a color gradient ranging from pink to black with increasing incubation times. (**B**) Pair distance distribution, P(r), as obtained at 1 mg mL$^{-1}$. (**C**) Kratky plot of the SAXS data obtained at 1 mg mL$^{-1}$. The same color code as in panel A was used.

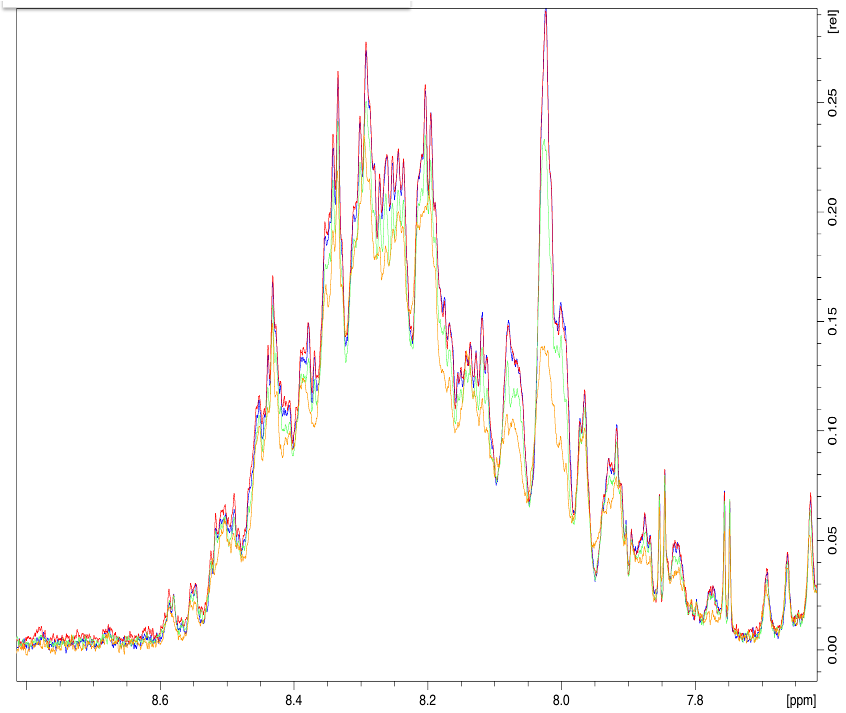


**Supplementary Figure S2.** 1D NMR spectra of a PNT3 sample at 100 µM after 0 (black), 8 (red), 27 (green) and 56 (yellow) hours of incubation at 37°C.

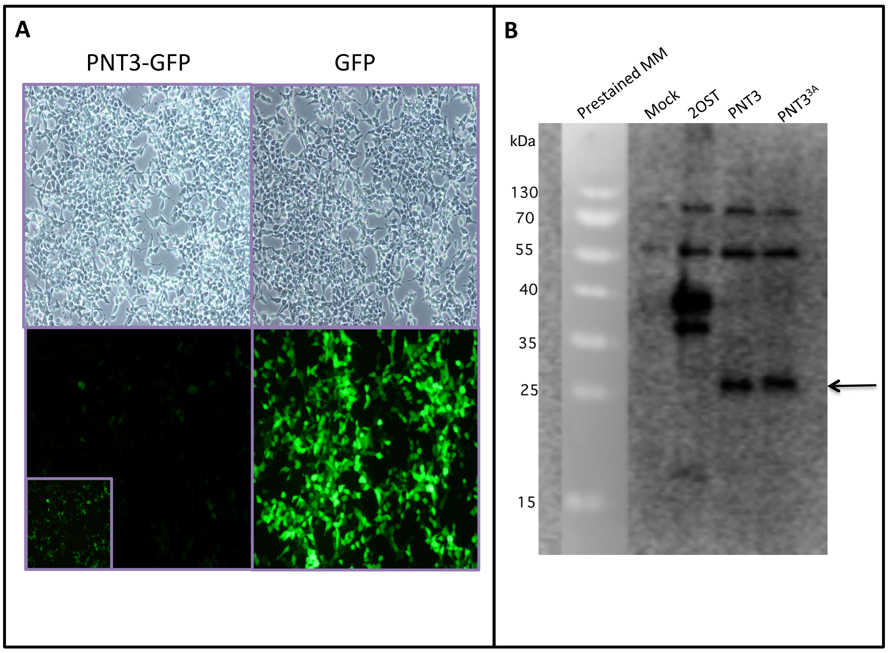


**Supplementary Figure S3. (A)** Non transfected HEK cells and transfected cells to express PNT3-GFP or GFP were observed under an epi-fluorescent microscope (magnification X100) with a contrast phase device (top images) and with a GFP filter (ex 488 nm, bottom images). Note the higher fluorescence of cells expressing GFP, reflecting a higher expression of GFP compared to PNT3-GFP. (**B**) Western Blot analysis of HEK293T cell lysates either non transfected (mock) or transfected to express an irrelevant control protein (2OST), PNT3 or PNT3$^{3A}$, using a Penta-His HRP conjugate. MM: molecular mass markers. The arrow indicates *wt* or mutated PNT3.

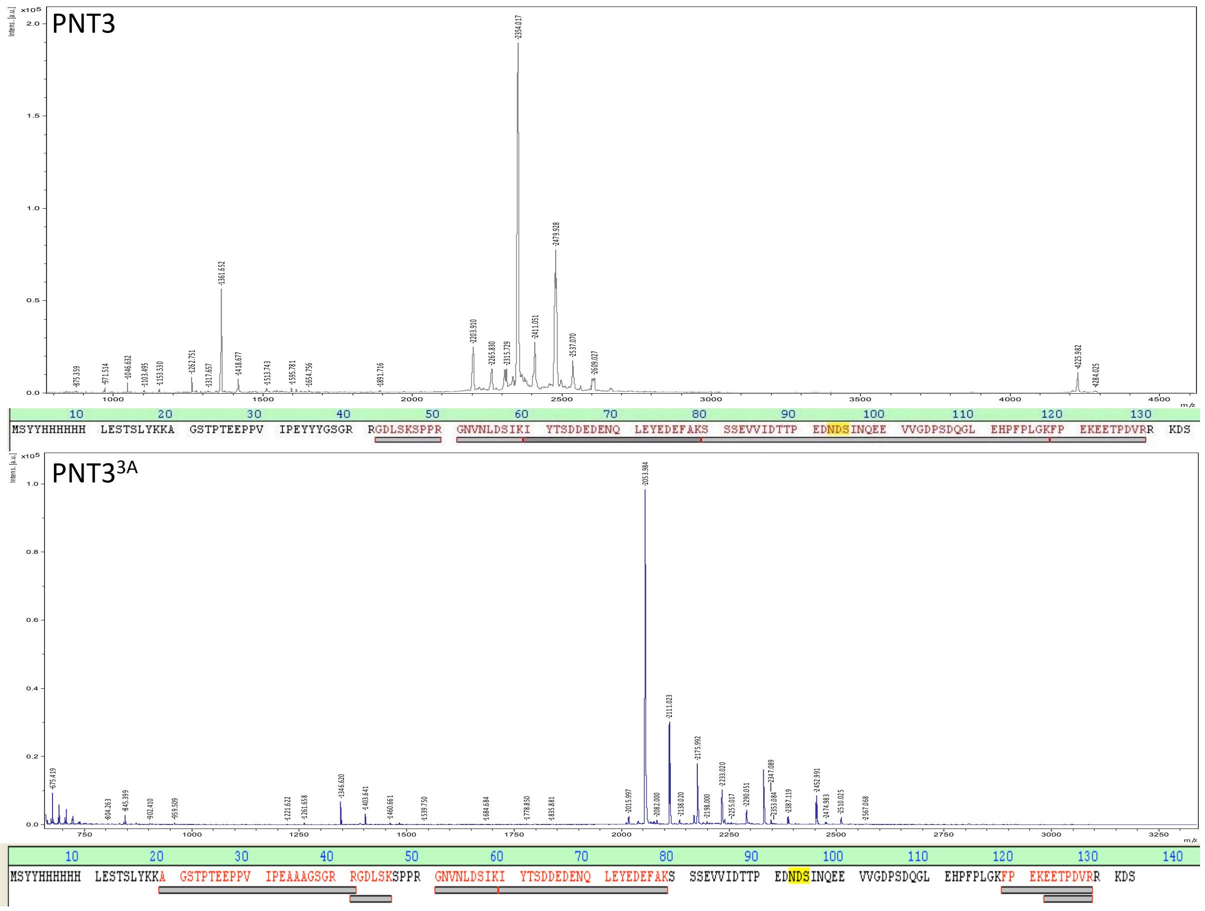


**Supplementary Figure S4.** Mass spectrometry analysis of PNT3 (**A**) and PNT3$^{3A}$ (**B**). The band of the protein was excised from a SDS acrylamide gel and then subjected to trypsin digestion. A coverage of 66% and of 50% of the sequence was obtained for PNT3 and for PNT3$^{3A}$, respectively. Peptides identified in the MS spectrum are highlighted by grey bars, with a color gradient ranging from light to dark grey with increasing peak intensities.

**Supplementary Table S1.** Molecular mass (MM), radius of gyration ($R_g$) and maximal intramolecular distance ($D_{max}$) at various time points of incubation at 37°C at two PNT3 concentrations.

| | **1 mg mL$^{-1}$** | | | **2 mg mL$^{-1}$** | | |
|---|---|---|---|---|---|---|
| Time (min) | MM | $R_g$ | $D_{max}$ | MM | $R_g$ | $D_{max}$ |
| 60 | 13.9 | 3.3 | 11.8 | 15.6 | 3.2 | 9.4 |
| 90 | 13.2 | 3.1 | 9.8 | 16.2 | 3.2 | 10.8 |
| 120 | 21.0 | 3.1 | 11.1 | 290.6 | 12.6 | 45.2 |
| 180 | 22.0 | 3.9 | 13.0 | 303.9 | 12.3 | 42.2 |
| 270 | 336.0 | 15.1 | 56.0 | 450.1 | 10.9 | 43.2 |
| 300 | 367.5 | 16.8 | 66.4 | 535.1 | 11.2 | 42.5 |
| 420 | 337.9 | 14.5 | 56.2 | 677.9 | 13.3 | 52.3 |
| 480 | 713.3 | 13.8 | 56.6 | 1235.5 | 13.3 | 52.6 |
| 630 | 1578.5 | 14.5 | 57.4 | 1565.5 | 13.9 | 52.9 |

**Supplementary Table S2.** Nucleotide sequence of the primers used to generate the various constructs.

| Primer name | Sequence (5'-3') | Name of the construct |
|---|---|---|
| PNT1-AttB1<br>PNT1-AttB2 | GGGGACAAGTTTGTACAAAAAAGCAGGCTTCGATAAACTGGATCTGGTTAACG<br>GGGGACCACTTTGTACAAGAAAGCTGGGTCTTATTACATCGGATCCAGCTGGATATC | PNT1-pDEST17 |
| PNT2-AttB1<br>PNT2-AttB2 | GGGGACAAGTTTGTACAAAAAAGCAGGCTTCGAAGATCCGGATGATATCCAG<br>GGGGACCACTTTGTACAAGAAAGCTGGGTCTTATTATTCCGGAATAACCGGCGGTTC | PNT2-pDEST17 |
| PNT3-AttB1<br>PNT3-AttB2 | GGGGACAAGTTTGTACAAAAAAGCAGGCTTCACCCCGACCGAAGAACCGCCG<br>GGGGACCACTTTGTACAAGAAAGCTGGGTCTTATTAGCTATCTTTACGACGCACATC | PNT3-pDEST17 |
| PNT4-AttB1<br>PNT4-AttB2 | GGGGACAAGTTTGTACAAAAAAGCAGGCTTCGAAGAAACCCCGGATGTGCGT<br>GGGGACCACTTTGTACAAGAAAGCTGGGTCTTATTATTTTTTGATCGGCATAATACGGC | PNT4-pDEST17 |
| B1HisNT3<br>NT3B2 | GTACAAAAAAGCAGGCTCGCATCACCACCATCACCATACCCCGACCGAAGAACCGCC<br>ACCACTTTGTACAAGAAAGCTGGGTcGCTATCTTTACGACGCACATC | His6-PNT3-pTH31 |
| HindNT3<br>NT3R<br>GFPqF<br>GFPqXhoI | GCGTTTAAACTTAAGCTTCCACCATGGCTAGCGATCAAACAAG<br>GGTGGCGACCGGTACCCATCG<br>CGATGGGTACCGGTCGCCACCATGGCTAGCAAAGGAGAAG<br>GGGCCCTCTAGACTCGAGTTATCAGTTGTACAGTTCATCC | His-PNT3-GFPq-pCDNA3.1+ |
| HindNT3<br>NT3XhoI | GCGTTTAAACTTAAGCTTCCACCATGGCTAGCGATCAAACAAG<br>TTAAACGGGCCCTCTAGACTCGAGTTATTAGGTGGCGACCGGTACCCATCG | His-PNT3-pCDNA3.1+ |
| attB1<br>R_3ala-pNT3<br>F_3ala-pNT3<br>attB2 | ACAAGTTTGTACAAAAAAGCAGGCT<br>TCGCCACGACGGCCGCTACCGGCTGCCGCTTCCGGAATAACCGGCGGTTCTTC<br>AACCGCCGGTTATTCCGGAAGCGGCAGCCGGTAGCGGCCGTCGTGGCGATCTG<br>ACCACTTTGTACAAGAAAGCTGGGT | mutPNT3-pDEST17 |
| HindNT3<br>R_3ala-pNT3<br>F_3ala-pNT3<br>NT3XhoI | GCGTTTAAACTTAAGCTTCCACCATGGCTAGCGATCAAACAAG<br>TCGCCACGACGGCCGCTACCGGCTGCCGCTTCCGGAATAACCGGCGGTTCTTC<br>AACCGCCGGTTATTCCGGAAGCGGCAGCCGGTAGCGGCCGTCGTGGCGATCTG<br>TTAAACGGGCCCTCTAGACTCGAGTTATTAGGTGGCGACCGGTACCCATCG | His-PNT3-3A-pCDNA3.1+ |
